# Predicting species and community responses to global change in Australian mountain ecosystems using structured expert judgement

**DOI:** 10.1101/2020.09.23.309377

**Authors:** James S. Camac, Kate D.L. Umbers, John W. Morgan, Sonya R. Geange, Anca Hanea, Rachel A. Slatyer, Keith L. McDougall, Susanna E. Venn, Peter A. Vesk, Ary A. Hoffmann, Adrienne B. Nicotra

## Abstract

Conservation managers are under increasing pressure to make decisions about the allocation of finite resources to protect biodiversity under a changing climate. However, the impacts of climate and global change drivers on species are outpacing our capacity to collect the empirical data necessary to inform these decisions. This is particularly the case in the Australian Alps which has already undergone recent changes in climate and experienced more frequent large-scale bushfires. In lieu of empirical data, we used a structured expert elicitation method (the IDEA protocol) to estimate the abundance and distribution of nine vegetation groups and 89 Australian alpine and subalpine species by the year 2050. Experts predicted that most alpine vegetation communities would decline in extent by 2050; only woodlands and heathlands were predicted to increase in extent. Predicted species-level responses for alpine plants and animals were highly variable and uncertain. In general, alpine plants spanned the range of possible responses, with some expected to increase, decrease or not change in cover. By contrast, almost all animal species were predicted to decline or not change in abundance or elevation range; more species with water-centric life-cycles were expected to decline in abundance than other species. In the face of rapid change and a paucity of data, the method and outcomes outlined here provide a pragmatic and coherent basis upon which to start informing conservation policy and management, although this approach does not diminish the importance of collecting long-term ecological data.

**Article Impact Statement:** Expert knowledge is used to quantify the adaptive capacity and thus, the risk posed by global change, to Australian mountain flora and fauna.

## Introduction

Alpine, subalpine and montane species are predicted to be negatively impacted by climate change. For the most part, this is because the climate envelope for many mountain species is expected to shrink and, in some regions, disappear entirely as a consequence of increased global temperatures (Halloy & Mark 2003; La Sorte & Jetz 2010; Freeman et al. 2018). While range contractions have already been observed in some mountain plants (Grabherr et al. 1994; Lenoir et al. 2008; Steinbauer et al. 2020) and animals (Freeman et al. 2018, Wilson et al. 2005), not all species are responding to climate change in the same way (Lenoir et al. 2010; Tingley et al. 2012; Gibson-Reinemer & Rahel 2015). What remains unclear is the capacity of mountain species to adapt (Hargreaves et al. 2014; Michalet et al. 2014; Normand et al. 2014; Louthan et al. 2015), and the characteristics that allow species to persist in the face of a changing climate (Fordham et al. 2012; Foden et al. 2018).

To understand the complexities and uncertainties of species responses to climate change, there have been several attempts to quantify adaptive capacity (Foden et al. 2013; Ofori et al. 2017; Gallagher et al. 2019). Adaptive capacity describes the ability of systems and organisms to persist and adjust to threats, to take advantage of opportunities, and/or to respond to change (Millenium Ecosystem Assessment 2005; IPCC 2014). Adaptive capacity confers resilience to perturbation, allowing ecological systems to reconfigure themselves with change (Holling 1973). In the context of alpine biota in Australia, adaptive capacity is the ability of species to maintain their often limited geographical distributions and population abundance when the climate and other factors are altered. While the underlying factors determining adaptive capacity encompass genetic and epigenetic variation, life history traits and phenotypic plasticity (Dawson et al. 2011; Ofori et al. 2017), little is known about which taxa have high adaptive capacity, how to quantify it, how it varies within and across related species, or how to manage populations in order to maximise it. As a consequence, data required to advise on the adaptive capacity of species are lacking.

Nonetheless, conservation practitioners and land managers are under increasing pressure to make decisions about the allocation of finite resources used to conserve biodiversity under climate change. Decisions are typically based on vulnerability assessments that incorporate exposure risk, species sensitivity, and adaptive capacity (Foden et al. 2013; Ofori et al. 2017; Foden et al. 2018). Until now, assessments of potential climate change impacts on species that cover multiple taxonomic groups have been based primarily on species distribution models (e.g. Thomas et al. 2004; Lawler et al. 2009; La Sorte & Jetz 2010). Incorporating species’ physiological, ecological and evolutionary characteristics, in conjunction with their predicted climate change exposure, will likely facilitate accurate identification of the species most at risk from climate change (Briscoe et al. 2020). However, these assessments focus on changes in species’ distribution or extent, their ‘climate space’, and the abiotic and biotic stresses that affect population ecology and physiology are not always fully represented in them (Guisan & Thuiller 2005; Geyer et al. 2011; Fordham et al. 2012). Further, the required data are rarely available for most species and the technical skill and time required to build and fit relevant models restrict their use to specialists (Briscoe et al. 2020). Given that the rate of climate change impacts has already outpaced our capacity to collect the required data to assess species empirically, it is important to utilise alternative methods that make use of existing expertise across taxa to estimate adaptive capacity and identify conservation priorities (Granger Morgan et al. 2001).

The need to predict how species will respond to climate change is particularly pertinent to the Australian alpine ecosystem which has a high level of endemism and a restricted geographic range (Venn et al. 2017). Since 1979, mean spring temperatures in the Australian Alps have risen by approximately 0.4 °C and annual precipitation has fallen by 6% (Wahren et al. 2013), with a consequent decline in snow pack depth (Sanchez-Bayo & Green 2013). Snow cover in Australia is now at its lowest in the past 2000 years (McGowan et al. 2018). These climatic changes correlate with changes in floristic structure, abundance and diversity (Wahren et al. 2013; Camac et al. 2015) and increases in fire frequency and severity (Camac et al. 2017; Zylstra 2018). Changes are expected to threaten the many locally adapted and endemic species, with cascading effects on biodiversity and ecosystem services such as carbon storage and water yield.

Here, we used a structured expert elicitation framework called the IDEA (“Investigate”, “Discuss”, “Estimate” and “Aggregate”) protocol (Hemming et al. 2018) to quantify changes in Australian alpine species’ future abundance in light of the many threats to their persistence. Structured expert elicitation provides a robust framework to estimate risk when data are either inadequate or lacking entirely (Hemming et al. 2018). While structured expert elicitation is increasingly being used in policy and management, few examples of its use exist in the ecological and conservation literature (Hemming et al. 2018). Expert elicitation quantitatively harnesses the local knowledge of biologists, conservation scientists, and natural resource managers to make predictions about critical but data-poor processes.

In this study, 37 experts (Table S1) estimated changes in the future abundance and/or distribution of nine Australian alpine plant communities, 60 alpine plant species and 29 mountain animal species. Expert knowledge provided insights into the species’ attributes and the biotic and abiotic factors that were expected to influence a species’ adaptive capacity. Using these expert elicited data, we:

1. quantified the direction and magnitude of change in cover/abundance/elevation range of Australian mountain plant communities as well as individual plant and animal species to climatic changes expected by 2050;
2. examined species attributes and biotic and abiotic factors that experts used when predicting changes in community and species abundances and how they compared to broad concepts about determinants of adaptive capacity, and;
3. examined how various measurable species attributes correlated with predicted changes in plant species abundance.

## Methods

### Study system

Australian high mountain ecosystems are restricted to south-eastern Australia, occupying an area ~ 11700 km^2^, or 0.15% of the continent. They are comparatively low in elevation, barely exceeding 2000 m a.s.l, ancient and mostly covered in soils. There is no nival zone or areas of permanent snow and some alpine areas of Tasmania even remain snow-free during the winter (Venn et al. 2017).

Australian mainland alpine ecosystems encompass several plant communities characterised by different species and growth forms (Kirkpatrick & Bridle 1999; Williams et al. 2006; Venn et al. 2017). Heathland predominates on relatively steep sheltered slopes where alpine humus soils are shallow (<0.3 m deep). The shrubs are 1–2 m tall, with a canopy cover typically exceeding 70%. Grassland/herbfield complexes occupy the more level ground on slopes and hollows, some of which may be subject to severe winds and frost, and where the alpine humus soils are deepest (generally up to 1 m). Short herbfields (i.e. snowpatch vegetation) occur on steep, leeward, south-to east-facing slopes where snow persists well into the spring or summer (Venn et al. 2017). Feldmark are an extremely rare ecosystem, existing only on exposed rocky ridges consisting of prostrate, hardy shrubs of the family Ericaceae. Wetland complexes consist of heathlands, bogs and fens and occupy valley bottoms, drainage lines and some stream banks and are typically waterlogged for at least one month per year. Wet tussock grasslands are regularly inundated with water or snowmelt, also at lower parts of the landscape. Woodlands are dominated by multi-stemmed, slow-growing trees (*Eucalyptus pauciflora*) and are typically snow-covered for at least one month each year.

The abundance and activity of the animals are regulated by the seasons (Green & Osborne 1994; Green & Stein 2015). The fauna consists of seasonal migrants and alpine specialists and is dominated by insects and other invertebrates (Green & Osborne 1994, Green & Slatyer 2020). Many species appear to be semelparous and require the snow pack to protect their overwintering eggs (e.g. *Kosciuscola* grasshoppers). Others, such as the *Monistria* grasshoppers, can overwinter as adults in the subnivial space by supercooling and thus have overlapping generations. Many Australian alpine insects exhibit iconic behaviour such as the long-distance migration of bogong moths (*Agrotis infusa*) (Warrant et al. 2016) or the striking startle display of the mountain katydid (*Acripeza reticulata*) (Umbers & Mappes 2015). The streams and wetlands support large alpine crayfish (*Euastacus spp*.), endemic earthworms (e.g. *Notoscolex montiskosciuskoi*), galaxiid fish, and several terrestrial-breeding frogs. The reptile diversity includes elapid snakes and many skink species. Most birds leave the alps in winter, returning to forage each summer. The only alpine endemic marsupial, the mountain pygmy possum (*Burramys parvus*), hibernates in boulder fields under the snow (Geiser & Broome 1991) while other mammals, such as wombats and echidnas, remain active throughout winter.

### Applying the IDEA protocol for structured expert elicitation

We utilised the IDEA protocol for structured elicitation of expert judgement (Hemming et al. 2018; Fig S1). This protocol involved: 1) recruiting a diverse group of experts to answer questions with probabilistic or quantitative responses; 2) discussing the questions (Table S2) and clarifying their meaning, and then providing private, individual best estimates and associated credible intervals, often using either a 3-point (i.e. best estimate, lower and upper limit; animal workshop) or 4-point (i.e. best estimate, lower and upper limit and confidence that the true value falls within those limits; plant workshop) elicitation method (Spiers-Bridge et al. 2010); 3) providing feedback on the experts’ estimates in relation to other experts; 4) discussing the results as a group, resolving different interpretations of the questions, sharing reasoning and evidence, and then providing a second and final private estimate, and; 5) aggregating experts’ final estimates mathematically, including exploration of performance based weighting schemes of aggregation (see also Supplemental Material).

The plant and animal expert elicitation projects were undertaken in July 2017 and November 2018, respectively. Because there is no accepted method to quantify or compare adaptive capacity across plants and animals, we developed questions based on estimates of percent cover for plants or abundance/elevation range for animals for the present day and in 2050. Experts (*n* = 22 for plants, *n* = 17 for animals, *n* = 2 shared between workshops; Table S1) were selected to represent a breadth of expertise in alpine botany, zoology and ecology in Australia. In the plant workshop, experts estimated the current (2017) and the 2050 cover of 60 plant species (Table S4), with 10 to 15 representative species in each of five dominant alpine vegetation communities. Furthermore, experts estimated the future landscape cover of nine alpine/subalpine vegetation community complexes based on an agreed 2017 baseline cover: feldmark (0.1%), snowpatch (1%), grassland/herbfield (25%), woodland (24%), heathland (35%), bog (5%), fen (4%) and wet tussock grassland (6%). For the plant elicitation, we assumed increases in temperature, decreases in precipitation (and less of that falling as snow, and fewer days of snow cover), and increased chance of fire. For the animal elicitation, we provided a specific climate scenario for the year 2050 (Table S3).

Expert-derived data is often aggregated in one of two ways, weighted or equally weighted. Our analysis focused on using equally weighted *best* estimates from experts. While expert uncertainty defined by their bounds and estimated confidence was collected in both workshops, it was not used in this analysis due to considerable variability in how experts interpreted, and thus, estimated their bounds (see Supplemental Material).

### Data Analysis

#### Calculation of summary statistics

We calculated the mean and 95% confidence intervals under both current and future scenarios for each species or plant community type. Various data transformations were required to estimate the mean and confidence limits because estimates were bounded (e.g. percent cover and abundance). For the plant percent cover data, individual expert best estimates were first logit transformed and then both mean and 95% confidence limits were estimated. Inverse logit transformations were then applied to each summary statistic to convert these estimates back to a proportional scale. As the animal abundance estimates were based on species-specific spatial scales, we first re-scaled expert estimates to a standard spatial scale (i.e. 100 m^2^). As some experts included zeros in their best estimates of abundance and elevation estimates, we applied a small constant (0.1) prior to log transforming the data. Means and 95% confidence limits were then calculated and back transformed to their original scale. Means and confidence limits for expert estimates of elevation range (maximum elevation minus minimum elevation) were calculated on the raw scale (i.e. not transformed prior to estimation). Comparison between ‘present’ and ‘future’ estimates was done using ‘inference by eye’ (Cumming & Finch 2005) by examining whether the 95% confidence intervals crossed the 1:1 line in plots of current vs future estimates. Finally, we used individual expert current and future best estimates to calculate the proportion of experts that indicated increase, decrease or no change.

To determine whether the change projected by the experts for alpine plants correlated with available data on species traits or environmental attributes, we calculated a proportional change in cover estimated by each expert (See Supplementary Material). Means and confidence intervals were then estimated and used to calculate the spearman rank correlations between this proxy of adaptive capacity and 1) a set of environmental measures derived from records in the Australian Virtual Herbarium and 2) plant functional trait data obtained from the experts’ published and unpublished data, as well as other published and online sources and, for a few species, field specimens were collected to supplement available data.

De-identified data and code used to produce figures 1–4 and Supplementary figures S2-S4 can be found at: https://github.com/jscamac/Alpine_Elicitation_Project.

**Fig 1.**
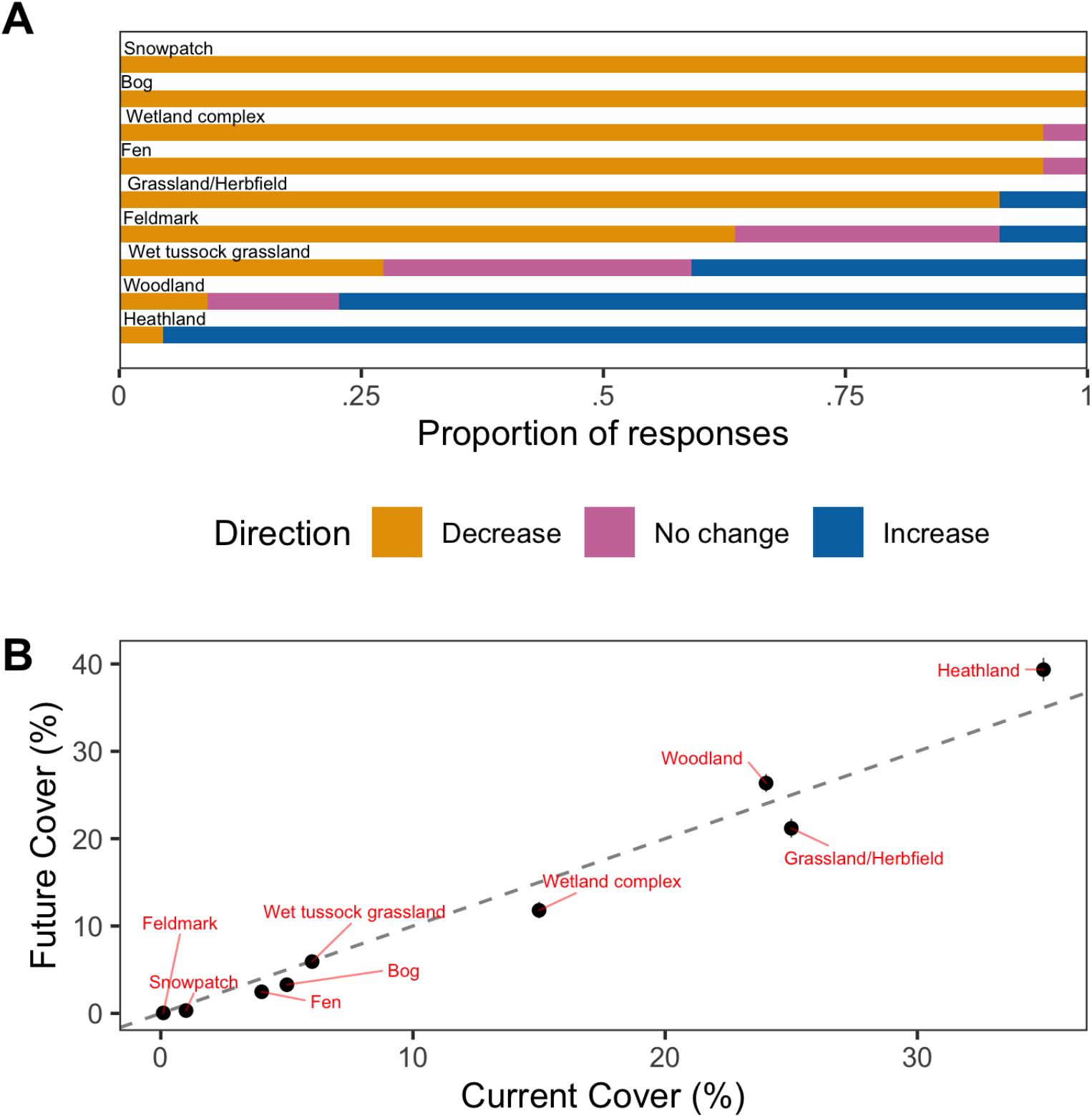
Nine Australian alpine plant community landscape cover predictions for 2050. A) The proportion of experts’ (*n* = 22) best estimates indicating a decline (orange), no change (pink) or increase (blue) in landscape cover between 2017 and 2050. B) Mean (± 95% confidence intervals) of expert best estimates of community landscape cover for 2050. Records below the dashed 1:1 line signify a decrease in cover, while those above the line signify an increase in cover. Assumed current landscape covers were agreed upon by experts: Feldmark (0.1%), Snowpatch (1%), Grassland/Herbfield (25%), Woodland (24%), Heathland (35%), Bog (5%), Fen (4%), Wet tussock grassland (6%).

**Fig 2.**
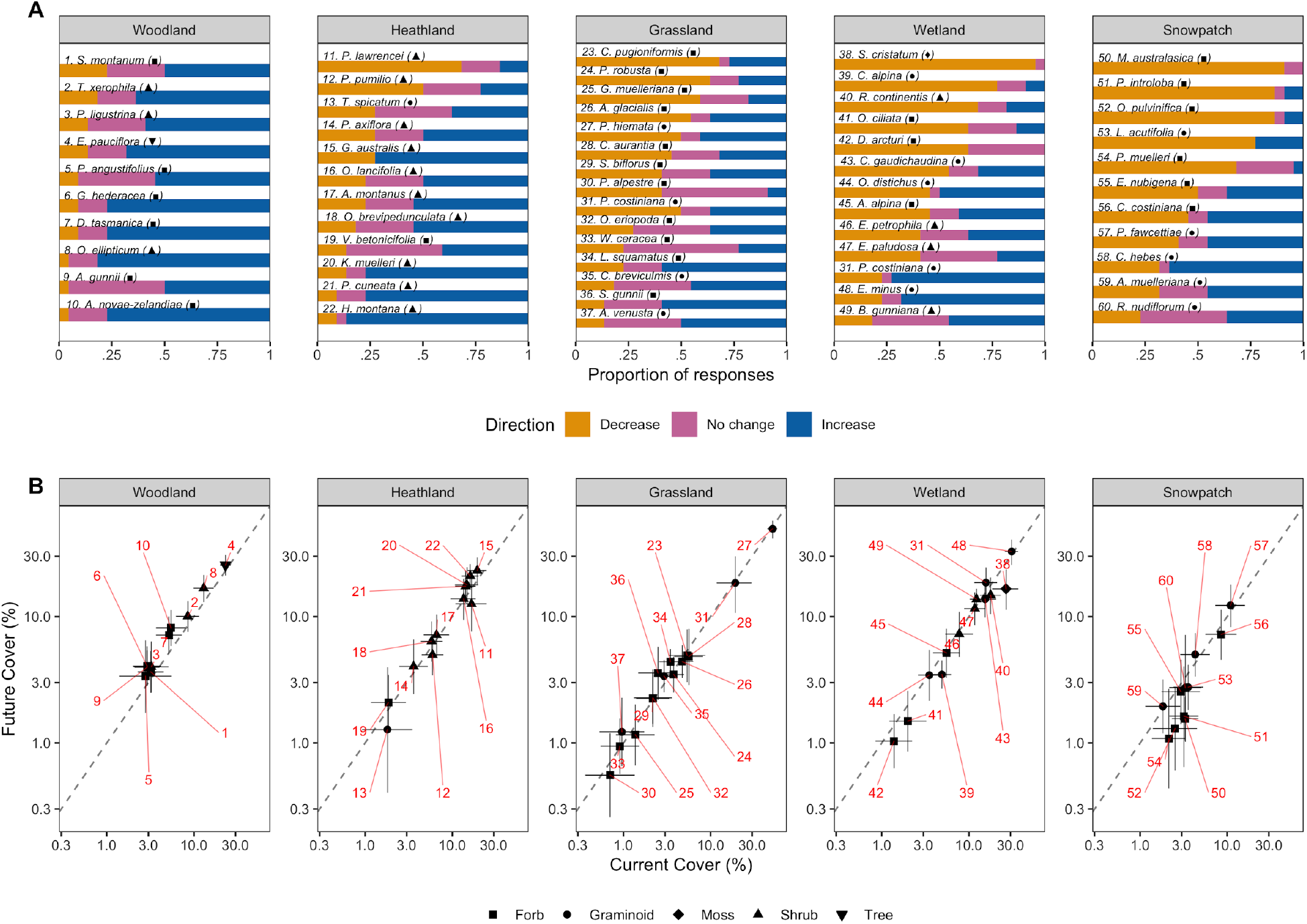
Sixty Australian alpine plants species cover predictions for 2017 and 2050. A) The proportion of experts’ (*n* = 22) best estimates indicating a decline (orange), no change (pink) or increase (blue) in cover between 2017 and 2050. B) Mean (± 95% confidence intervals) of expert best estimates of species cover for 2017 and 2050. Records above the dashed 1:1 line signify a decrease in cover, while those above the line signify an increase in cover. Species have been grouped by the community type they most commonly occur in. Numbers signify species ID.

**Fig 3.**
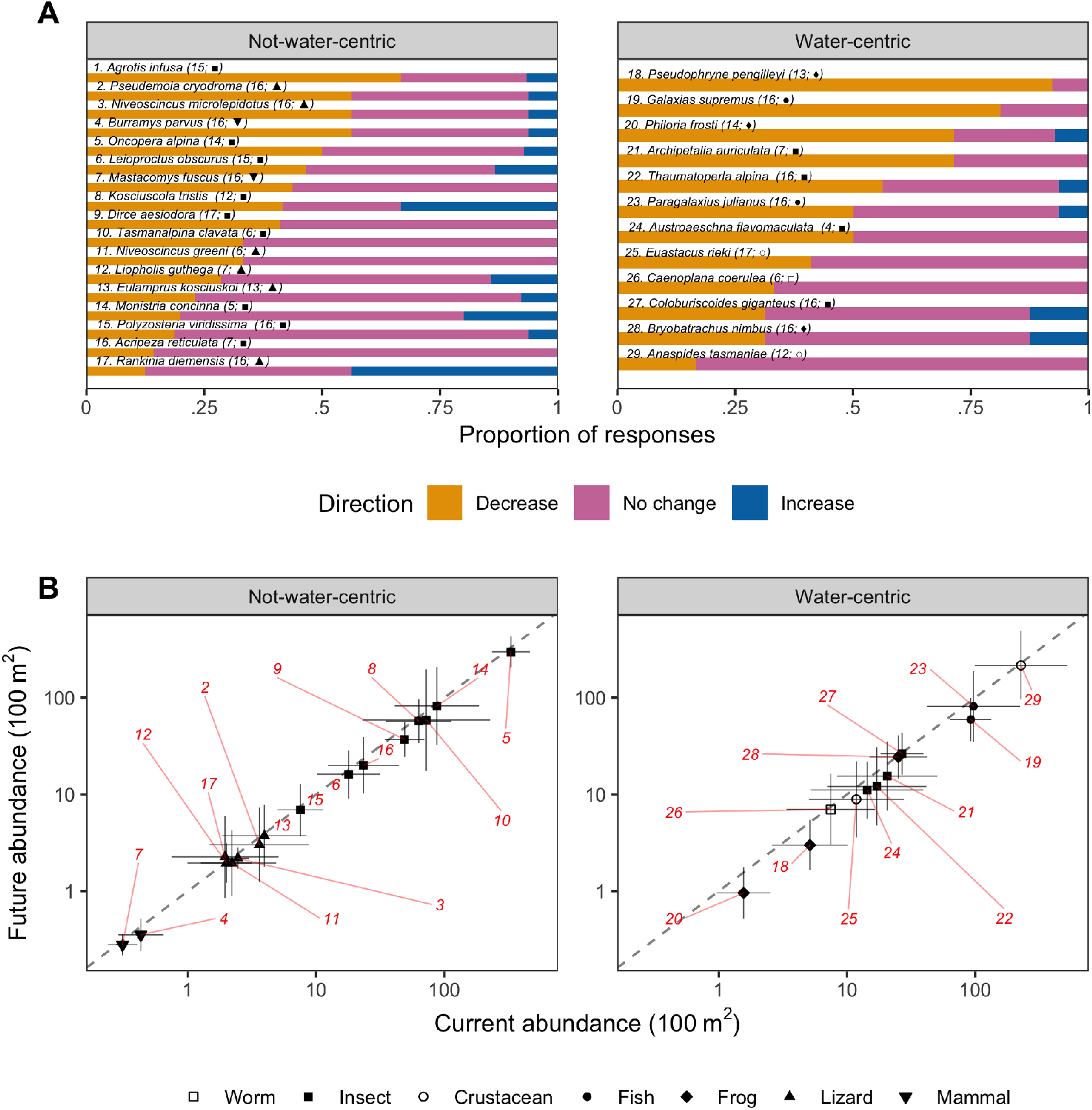
Twenty-nine Australian alpine animal species’ abundance predictions for 2018 and 2050. A) The proportion of experts best estimate indicating a decline (orange), no change (pink) or increase (blue) in cover in 2018 and 2050. B) Mean (± 95% confidence intervals) of expert best estimates of species abundance for 2018 and 2050. Records above the dashed 1:1 line signify a decrease in abundance, while those above the line signify an increase in abundance. Species are grouped by degree of dependency on water to complete their life-cycle as water-centric and non-water-centric. Numbers signify species ID. Numbers in parentheses in panel (A) represent the number of experts who provided estimates (Maximum = 17). Symbols represent higher taxon. Note: the bogong moth (*A. infusa*) has been omitted from panel B as its abundance estimates were multiple orders of magnitude higher than other species.

**Fig 4.**
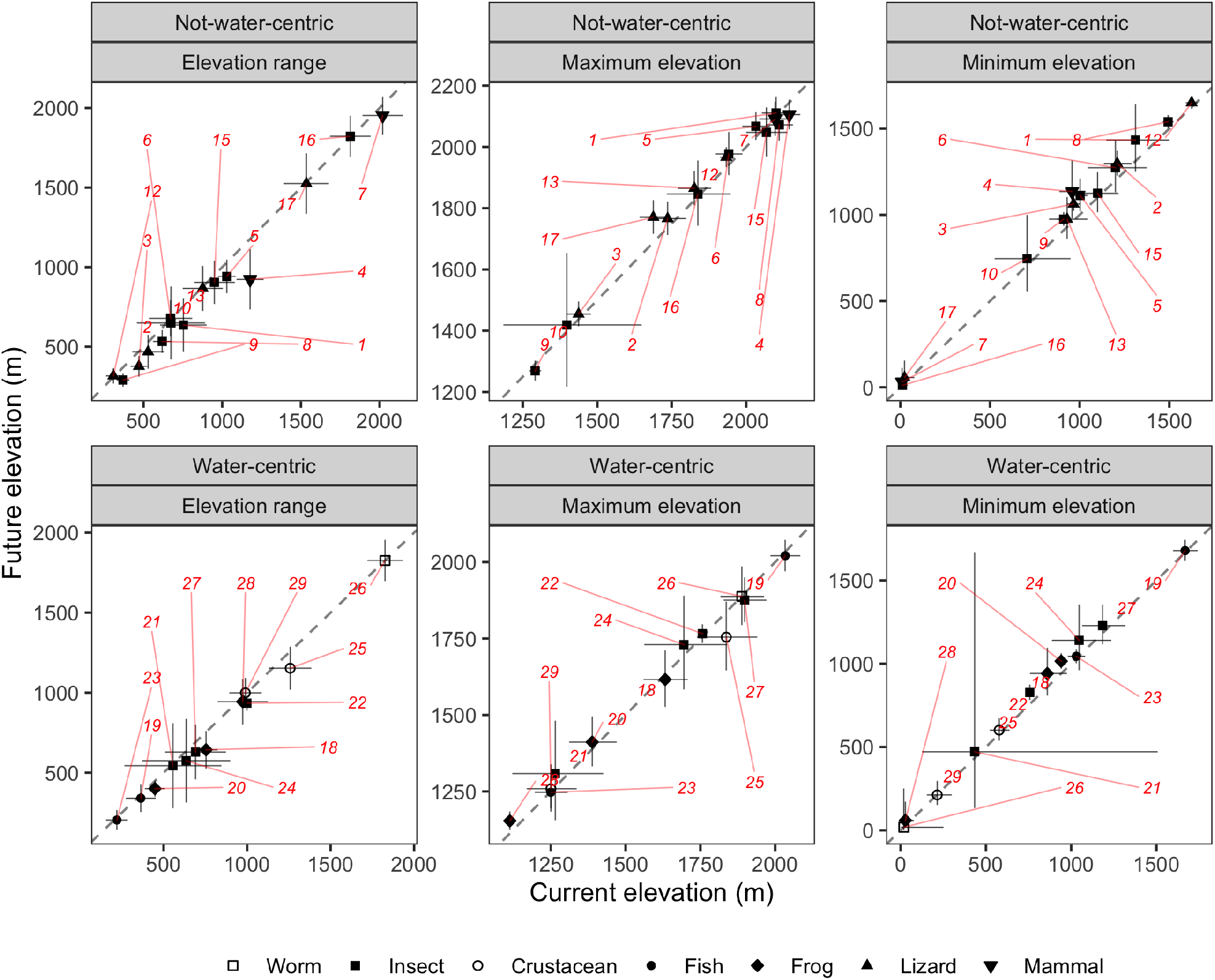
Australian alpine fauna species mean (± 95% confidence intervals) elevation range (left panels); maximum elevation (center panels) and minimum elevation (right panels) predictions for 2018 and 2050. Records below the dashed 1:1 line signify a decrease, while those above the line signify an increase. Species are grouped by degree of dependency on water to complete their life-cycle, as water-centric and non-water-centric Numbers signify species ID (see Fig 3A). Symbols represent taxon class.

## Results

### Predicted change in cover of Australian mountain vegetation types

Most of alpine vegetation communities were predicted by the majority of experts to decline in extent (i.e. total cover in the landscape) with global change by 2050 (i.e. snowpatch, bog, fen, wetland complex, grassland/herbfield). All experts predicted that snowpatch and bog communities will decrease by 2050, whereas most experts predicted heathlands and woodlands would increase in extent (Fig 1A). There was more uncertainty among experts about the future of wet tussock grasslands and feldmark communities (Fig 1A). Communities that are currently restricted in extent across the Australian alpine landscape (<5% extent) were predicted to be the ones most likely to decline (Fig 1B), but some of the more extensive communities (i.e. wetland complex, grassland/herbfield, which currently occupy ~25% of the landscape) were also predicted to decline in extent (Fig 1B).

### Direction and magnitude of change in cover for individual plant species

Within each plant community, experts predicted that the individual species’ responses to global change would vary (Fig 2). Some species, such as the snowpatch forb *Montia australasica* (#50 in Fig 2) and the wetland moss *Sphagnum cristatum* (#38), were almost unanimously predicted to decline in cover over time (Fig 2A). For other species, such as the subalpine heathland shrub *Hovea montana* (#22), experts predicted increases in cover (Fig 2A), although the magnitude of increase was small (Fig 2B). For most alpine plant species, there was much uncertainty about their future cover relative to current cover. The snowpatch graminoid *Rytidosperma nudiflorum* (#60), the wetland shrub *Baeckea gunniana* (#49), the grassland forb *Oreomyrrhis eriopoda* (#32), the heathland shrub *Acrothamnus montanus* (#17), the woodland forb *Stylidium montanum* (#1) and even the grassland structural dominant *Poa hiemata* (#27) were, according to experts, equally likely to show increases, decreases, or no change in cover (Fig 2B). This is reflected in the high uncertainty seen in future cover estimates (i.e. vertical error bars) for these species (Fig 2B).

Across all plant species, growth form was found to be relatively important in explaining expert judgements of species’ adaptive capacity (Fig 2A). Woody plants (shrubs and one tree) were typically predicted to have higher adaptive capacity (i.e. show increases or no change in cover) relative to forbs and graminoids (Fig 2).

In general, plant species with current high cover in herbaceous communities (e.g. snow patches, grasslands and wetlands) were not predicted to become more dominant with climate change. Experts were uncertain about the future cover of many of these current high-cover herbaceous species (Fig 2). For example, the graminoids *Poa costiniana* (#31, grasslands), *Poa fawcettiae* (#57, snowpatches) and the forb *Celmisia costiniana* (#56, snowpatches) were predicted by experts to either increase or decrease in cover in roughly equal numbers (Fig 2A). By contrast, in communities dominated by woody plants (heathlands, woodland), species with current high cover were predicted to increase their cover into the future (Fig 2B, e.g. *Hovea montana #22, Oxylobium ellipticum #8*).

### Direction and magnitude of change in abundance and elevation range for individual animal species

Animal expert predictions showed considerable variability in responses to global change (Fig 3). For nearly half the species (*n* = 13), the majority of experts predicted a decline in abundance (Fig 3A). The majority of experts suggested the Northern Corroboree Frog (*Pseudophryne pengellyi*, #18), the Baw Baw Frog (*Philoria frosti*, #20), the Kosciuszko Galaxis fish (*Galaxias supremus*, #19) and the Bogong Moth (*Agrotis infusa*, #1) would decline by 2050 (Fig 3A). For most of the remaining species, the majority of experts predicted no change in abundance. For example, most experts suggested that the abundance of the Mountain Katydid (*Acripeza reticulata*, #16) and the Mountain Shrimp (*Anaspides tasmaniae*, #29) will not change by 2050 (Fig 3A). There was no species for which the majority of experts predicted an increase in abundance, but a notable proportion of experts predicted an increase in the abundance of the Thermocolour Grasshopper (*Kosciuscola tristis* #8). Experts were split equally between ‘increase’ and ‘no change’ for the Mountain Dragon (*Rankinia diemensis*, #17) and split equally between ‘decrease’ and ‘no change’ for the Alpine Darner (*Austroaeschna flavomaculata*, #28) (Fig 3A).

Examining the magnitude of change in abundance (Fig 3B), many species were predicted to decline by 2050, although in almost all cases these changes were small and uncertain (i.e. confidence limits cross the 1:1 line). The exceptions to this were the Mountain Dragon (*Rankinia diemensis, #17*) which is predicted to marginally increase — although this is uncertain — and both the Northern Corroboree Frog (*Psuedophryne pengellyi, #18*) and the Baw Baw Frog (*Philoria frosti, #20*), which are predicted to likely decrease in abundance. Examining species responses across water-centric and non-water-centric life histories revealed that, on average, non-water-centric species were expected not to change in abundance, while water-centric species were more likely to decline.

With uncertainty, the minimum elevation limits of fauna distributions were predicted to shift upslope for 24 of 29 species (Fig 4; right panels). The Mountain Pygmy Possum (*Burramys parvus*, #4) had the largest predicted change in minimum elevation range-limit, expected to move up more than 150 m. The Alpine Cool Skink (*Niveoscincus microlepidotus* #3), Alpine Bog Skink (*Pseudemoia cryodroma*, #2) and Alpine Plaster Bee (*Leioproctus obscurus*, #6) also show substantial departures from no change. No change in minimum elevation was predicted for the two species whose distributions, while predominantly contained within mountain regions, extend to sea level – the Blue Planarian (*Caenoplana coerulea*, #26) and the Mountain Katydid (*Acripeza reticulata*, #16). The maximum elevation limits were predicted to increase for 16 species (range 8-80 m) and decrease for 11 species (range 1-80 m). Uncertainty encapsulated the 1:1 line for most species, but distinct increases in maximum elevation were predicted for the Mountain Dragon (*Rankinia diemensis*, #17). A conspicuous, but uncertain, reduction in maximum elevation was estimated for the alpine crayfish (*Euastacus reiki*, #25). For most species (*n* = 23), the total elevation range occupied was predicted to shrink as a result of upward shifts at low elevation limits. Increases in elevational range were predicted for four species and only one species - the Blue Planarian (*C. coerulea*, #26) - was predicted to show no change in elevational range by 2050. The largest declines in species elevational range were predicted for the Mountain Pygmy Possum (*Burramys parvus, #4*, ~250 m reduction), the Northern Corroboree Frog (*P. pengilleyi*, #18, ~110 m reduction) and the Alpine Crayfish (*Euastacus rieki*, #25, ~105 m reduction).

### Expert opinion on drivers of adaptive capacity

In the initial surveys, prior to the workshops, both plant and animal experts nominated genetic variability and phenotypic plasticity as key determinants of adaptive capacity, with fecundity, lifespan, and dispersal also considered important. However, notes and comments compiled during the elicitation process suggested that experts referred more often to environmental and biotic attributes when considering drivers of change in cover/abundance for specific organisms. Climate niche-breadth, disturbance regimes (e.g. fire, frost events) and species interactions, including competitive ability in the face of native (e.g. shrubs and trees) or exotic species encroachment (e.g. Horses, deer, weeds), vulnerability to diseases (e.g. *Phytophthora cinnamoni*) and a dependence on other species (e.g. grazers, pollinators), dominated discussions about potential drivers of future change in alpine species abundance and/or distribution.

### Correlations of plant species attributes with expert predictions

The projected magnitude of change in cover of plant species was correlated with environmental (Figure S2) and species range attributes (Figures S3 & S4). Adaptive capacity was most negatively correlated with species’ minimum elevation (r = −0.561) and most positively correlated with mean annual temperature range (r = 0.466), elevation range (r = 0.561) and area of occupancy (r = 0.43), noting that these three variables are themselves highly correlated with each other. We found that our measure of adaptive capacity was not strongly correlated with the continuous species traits such as mean height (r = 0.286), leaf area (r = −0.061), specific leaf area (r = −0.05), diaspore mass (r = 0.202) or dispersal distance (r = 0.342).

## Discussion

Conservation managers are increasingly required to make decisions about the allocation of finite resources to protect biodiversity under changing climate and disturbance regimes. Climate change impacts, however, are outpacing our capacity to collect data to assess individual risk empirically to inform resource allocation. A pragmatic alternative approach is to utilise expertise across taxa to produce timely estimates of conservation risk (Granger Morgan et al. 2001; Burgman et al. 2011a; Martin et al. 2012). Experts’ acquired experience allows them to provide valuable, nuanced insight into predictions about the future given a particular scenario. Our study has demonstrated the feasibility of a structured expert elicitation process for identifying the potential for adaptive capacity in Australian alpine plant communities, and individual animal and plant species. Adaptive capacity is the ability of systems and organisms to respond to consequences of change (IPCC 2014) and important for ecosystems undergoing rapid and substantial climate change such as alpine ecosystems (Steinbauer et al. 2018), tropical forests (Gallagher et al. 2019) and coral reefs (Silverstein et al. 2012). We identified that some alpine species and communities are likely to be more vulnerable to global change by 2050 than others. Our exercise also identified species for which experts are equivocal and thus, targets for further investigation.

Expert judgement identified that the adaptive capacity of Australian alpine biota in the face of global change is, not surprisingly, likely to be species-specific. Here, the adaptive capacity estimates encompassed more than just species’ responses to climate change; they also included structured consideration of all issues identified by experts such as a species’ response to fire, invasive species, predation and interspecific competition. While this may seem self-evident, it is the first time that multiple species and communities in alpine Australia have been simultaneously assessed for their adaptive capacity and it provides a defendable basis for targeting monitoring of vulnerable species and communities, as well as the development of potential mitigation strategies for at-risk species. When given a plausible 2050 climate change scenario, incorporating the assumption that an extensive bushfire would occur during this period (which subsequently happened in early 2020; Nolan et al. 2020), adaptive capacity was predicted to be lower in herbaceous plants relative to woody plants, and lower in water-centric animals relative to non-water-centric species. Adaptive capacity was not strongly correlated to quantitative plant traits such as specific leaf area or diaspore mass. This is perhaps unsurprising as such traits are thought to act on individual demographic rates (e.g. mortality, growth, fecundity), which themselves trade-off against one another. By contrast, adaptive capacity (i.e. proportional cover change) is the outcome of the amalgamation of multiple such trade-offs – thus diminishing possible correlations with individual traits. Moreover, the amount of inter-specific variation explained by traits typically assumed to be strongly linked to demographic rates (e.g. wood density and tree mortality) have been shown to be small (e.g. Camac et al. 2018). Unlike correlative species distribution models which rely only on climate data and species occurrence data, experts undertaking structured judgements inherently consider physiological, ecological and evolutionary characteristics of species, as well as how those species might interact (or re-assemble) in novel assemblages, and how disturbance (from fire in our case) may affect their responses.

We found that experts came into the elicitation process with perceptions of key environmental and biotic drivers of species responses to global change but, after discussion with other experts, they refined these drivers. Prior to the elicitation process, experts emphasized characteristics of the focal species as being the most important predictors of their response to global change (e.g. genetic variability, phenotypic plasticity, fecundity, lifespan, dispersal). During discussion, experts shifted their thinking to include both biotic and environmental drivers as being of importance to predicting alpine biota response to global change (e.g. competitive ability, mutualisms, niche breadth). This shows the value of using a structured elicitation method relative to informal elicitation approaches (Krueger et al. 2012).

As might be expected, ‘rare’ species - defined by animal abundance (or elevational range) or plant cover - were typically predicted to become rarer with global change. Small population size and restricted habitat breadth are likely key reasons for such thinking amongst experts (Williams et al. 2015; Cotto et al. 2017; Kobiv 2017). Terrestrial ectotherms (insects, reptiles, frogs), for example, are likely to face increased periods of heat stress (Hoffmann et al. 2013), while drought and declining snow cover duration make many plants and water-centric animals vulnerable (Wipf et al. 2009; Griffin & Hoffmann 2012; Williams et al. 2015). For many animals, experts predicted that species with the narrowest elevational range on mountains (such as the Mountain Pygmy Possum) are most likely to further contract. Such processes are already occurring in mountain landscapes, with lower limit upward shifts in species having already been reported (Pauli et al. 2007; Freeman et al. 2018; Rumpf et al. 2019).

Unexpectedly, experts were uncertain about the future abundance/cover of some ‘common’ species. While some structural dominants in plant communities are forecast to be either likely ‘winners’ (e.g. shrubs such as *Hovea montana, Grevillea australis, Prostanthera cuneata*) or ‘losers’ under global change (e.g. the moss *Sphagnum cristatum* in alpine wetland bogs), which is in broad agreement with other studies (e.g. Williams et al. 2015; Camac et al. 2017), there was less agreement about others. *Poa hiemata*, a dominant and potentially long-lived tussock grass of alpine grasslands and herbfields, had uncertain adaptive capacity according to experts. We suspect that experts varied in the emphasis they placed on a long adult lifespan in limiting the adaptive capacity of local populations, with longevity buffering individual persistence in unsuitable sites at least in the short-term (Cotto et al. 2017) but slowing evolutionary rates. Alternatively, experts were potentially weighting disturbance impacts, interspecific competition and climate sensitivity very differently (Granger Morgan et al. 2001). Given such species are functionally important, provide most of the community biomass (both above- and below-ground), structure habitat for fauna, and provide ecosystem services such as erosion control (i.e. they act as ‘foundation species’, Ellison & Degrassi 2017), understanding the autecology and dynamics of dominant species in response to global change drivers appears to be a key research need. Indeed, the uncertainty around common species responses highlights that long-term cover/abundance trends need to be quantified if future ecosystem stability is to be understood, a call that has been made repeatedly in the literature (Smith & Knapp 2003; Gaston & Fuller 2007; Gaston 2011; Smith et al. 2020). Monitoring species’ local abundance may therefore better inform species’ extinction risks in alpine areas under global change than monitoring their range (Cotto et al. 2017).

Overall, the change in cover of plant species, or elevational range and abundance change for animals, were estimated to be modest despite some climatic effects already becoming evident in Australia’s alpine biota (e.g. Camac et al. 2017; Hoffmann et al. 2019); estimates for cover change in plant communities were more pronounced. This may reflect that scientific experts are typically conservative when estimating the future (Oppenheimer et al. 2019). Experts also likely view biotic response to global change as a time-lagged process (i.e. ‘disequilibrium dynamics’, Svenning & Sandel 2013). Lags occur because of the limited ability of species to disperse to new areas (Morgan & Venn 2017; Alexander et al. 2018), establishment limitations following their arrival (Graae et al. 2011; HilleRisLambers et al. 2013; Camac et al. 2017), and the extinction debt of resident species (Dullinger et al. 2012). By forecasting only to 2050, experts have indicated that many longer-lived species will potentially persist through the initial ongoing change, but their capacity to do so beyond this is not assured. Lastly, biologists may find it difficult to estimate the rate of change. Most models of global change impacts are based on short-term experiments and have typically focused on differences or ratios of state variables (e.g. control vs manipulated groups). While these models are useful for inferring the direction of impacts (which implicitly inform expert views), they often do not provide information on the rate of change, the fundamental process needed to accurately forecast the magnitude of change (Camac et al. 2015; Morgan et al. 2016).

### Applicability of IDEA methodology to ecological problems

The IDEA protocol has been tested in a variety of application areas (Speirs-Bridge et al. 2010; Burgman et al. 2011a; McBride et al. 2012; Wintle et al. 2012, Hanea et al. 2016) and these tests consistently confirmed the value of using a diverse group of experts, of giving experts the opportunity to cross examine the estimates of their peers, and of reducing ambiguity through discussion. In our elicitations, we speculate that experts revised their initial estimates if they (i) had no direct knowledge of the species themselves but were guided by the discussion, (ii) aligned responses to those of a taxon specialist, or (iii) adjusted their values based upon a particular line of reasoning they found convincing during the discussion. Most validation studies found that when experts revise their estimates, they do so in the direction of the “truth” (e.g. Burgman et al. 2011b; Hanea et al. 2018).

One difficulty in using this methodology was revealed at both workshops - the capacity of the participants to undertake this particular kind of statistical estimation. Gigerenzer & Edwards (2003) and many others (e.g. Low Choy et al. 2009) have previously documented the difficulties experts have when communicating knowledge in numbers and probabilities. We attempted a four point elicitation with the plant experts for each species (1. lowest plausible value, 2. highest plausible value, 3. best estimate and 4. confidence that the truth falls between their lower and upper limits), and revised this down to a three point elicitation for the animal experts (by omitting the confidence estimate, and fixing the upper and lower limits to correspond to a central 90% credible interval). While experts were comfortable in providing best estimates, there was inconsistency (indeed confusion) about interpreting and estimating bounds and confidence - even after conducting a brief workshop outlining how to do it. For these reasons, our analysis focused on using each expert’s best estimates and not their estimated uncertainty defined by bounds and estimated confidence. Potentially valuable information about the confidence in estimates was therefore lost during the elicitation process. However, the IDEA protocol strives to elicit improved best estimates by eliciting bounds first. Even if the bounds are not used as a measure of the expert’s uncertainty, the counterfactual thinking needed prior to eliciting the best estimates improves the latter. We feel that the ‘best estimate’ of cover or abundance is useful for forecasting the direction and magnitude of change expected by experts under a given global change scenario. Moreover, we believe that involving a mechanism for discussing and revising estimates (through the IDEA protocol) provides robust insights into these potential changes.

### Management Implications

The adaptive capacity framework we used to elicit expert opinions about how alpine species and communities may respond to global change currently exists as a framework of “exposure risk” to change based on current state and predicted future state (i.e. our species prediction biplots). Our experts, through their judgment, implicitly accounted for multiple drivers of change in mountain ecosystems (e.g. rising temperatures, biotic interactions, feral animals, fire) but did so assuming no mitigation by management occurred. Using this approach, experts predicted that several plant (e.g. *Sphagnum cristatum*) and animal species (e.g. Baw Baw Frog *Philoria frosti*, Northern Corroboree Frog *Pseudophryne pengellyi*, and Mountain Pygmy Possum *Burrymus parvus*) appear very vulnerable to the changes in alpine areas that are predicted to occur by 2050.

If the value of the framework is to identify the species that are most vulnerable to global change (i.e. the species with limited adaptive capacity), then it becomes important to consider our capacity to influence adaptive capacity into the future through management intervention. This will be of most relevance to land managers and conservation biologists who want to reduce the risk of species extinction. We believe this will be critical to operationalise the expert judgment outcomes reported here. Having identified in our biplots which species have lower adaptive capacity, managers may begin to ask: how might we buffer them against climate change? Or, how can we improve the resilience of alpine species? There are many management actions that can reduce threats and these are already part of a land manager’s current arsenal such as removing feral animals and weeds, protecting vulnerable communities from fire and assisted migration.

If management actions could improve the adaptive capacity of alpine species, and these actions could be ranked for their efficacy to achieve such aims, then the expert judgements we have elicited in this study can be used to inform prioritisation for conservation actions in regions such as the Australian Alps. Hence, not only can we use a species’ adaptive capacity as a means to rank species in need of mitigation action, but we could identify the species most likely to respond to management interventions. Indeed, such an approach may even identify that, for some species, there is nothing that we can practically do to change their adaptive capacity. In such cases, it may be that options such as *ex situ* conservation strategies (such as seed banking, captive breeding) need to be implemented.

In an era of rapid change, conservation practitioners and land managers do not have the privilege of time to wait for additional data and knowledge to be accrued to inform their decisions. They must utilise information currently at hand to prioritise conservation efforts so that species losses may be mitigated. We believe the method and outcomes outlined here can provide a pragmatic and coherent basis for integrating available expert knowledge to quantify adaptive capacity and perhaps help mitigate the overwhelming risk posed by global change to the long-term persistence of Australian alpine species.

## Acknowledgements

This study was supported by funding from the National Climate Change Adaptation Research Facility National Adaptation Network for Natural Ecosystems (vegetation) and the Centre for Biodiversity Analysis, ANU and the NSW Dept. of Industry Conference Support Program (animals). Sandra Lavorel, Mel Schroder and Libby Rumpff helped refine our study scope and questions. We thank all experts who participated in the structured elicitation workshops. We also thank Linda Broome, Nick Clemann, Elaine Thomas and Phil Zylstra, who provided participants with critical information that was used to inform their estimates. Lastly, we thank Nola Umbers for taking on caring responsibilities for KU. The flora elicitation workshop was approved by the Human Ethics Committee of La Trobe University (Project Number: S17-069). The fauna elicitation workshop was approved by the Human Ethics Committee of Western Sydney University (Project Number: H12680).

## Literature Cited

Alexander JM, Chalmandrier L, Lenoir J, Burgess TI, Essl F, Haider S, Kueffer C, McDougall K, Milbau A, Nuñez MA, Pauchard A, Rabitsch W, Rew LJ, Sanders NJ, Pellissier L. 2018. Lags in the response of mountain plant communities to climate change. Global Change Biology 24: 563–579.

Briscoe NJ, Elith J, Salguero-Gómez R, Lahoz-Monfort JJ, Camac JS, Giljohann KM, Holden MH, Hradsky BA, Kearney MR, McMahon SM, Phillips BL, Regan TJ, Rhodes JR, Vesk PA, Wintle BA, Yen JD, Guillera-Arroita G. 2019. Forecasting species range dynamics with process-explicit models: matching methods to applications. Ecology Letters 22: 1940–1956.

Burgman M, Carr A, Godden L, Gregory R, McBride M, Flander L, Maguire L. 2011a. Redefining expertise and improving ecological judgment. Conservation Letters 4: 81–87.

Burgman MA, McBride M, Ashton R, Speirs-Bridge A, Flander L, Wintle B, Fidler F, Rumpff L, Twardy C. 2011b. Expert status and performance. PLoSOne 6: 1–7.

Camac JS, Williams RJ, Wahren C-H, Jarrad F, Hoffmann AA, Vesk PA. 2015. Modeling rates of life form cover change in burned and unburned alpine heathland subject to experimental warming. Oecologia 178: 615–628.

Camac JS, Williams RJ, Wahren C-H, Hoffmann AA, Vesk PA. 2017. Climatic warming strengthens a positive feedback between alpine shrubs and fire. Global Change Biology 23: 3249–3258.

Camac JS, Condit R, FitzJohn RG, McCalman L, Steinberg D, Westoby M, Wright SJ, Falster DS. 2018. Partitioning mortality into growth-dependent and growth-independent hazards across 203 tropical tree species. Proceedings of the National Academy of Sciences 115: 12459–12464.

Cotto O, Wessely J, Georges D, Klonner G, Schmid M, Dullinger S, Thuiller W, Guillaume F. 2017. A dynamic eco-evolutionary model predicts slow response of alpine plants to climate warming. Nature Communications 8: 15399.

Cumming G, Finch S. 2005. Inference by eye: confidence intervals and how to read pictures of data. American Psychologist 60: 170–80.

Dawson TP, Jackson ST, House JI, Prentice IC, Mace GM. 2011. Beyond predictions: biodiversity conservation in a changing climate. Science 332: 53–58.

Dullinger S, Gattringer A, Thuiller W, Moser D, Zimmermann NE, Guisan A, Willner W, Plutzar C, Leitner M, Mang T, Caccianiga M, Dirnbock T, Ertl S, Fischer A, Lenoir J, Svenning J-C, Psomas A, Schmatz DR, Silc U, Vittoz P, Hulber K. 2012. Extinction debt of high-mountain plants under twenty-first-century climate change. Nature Climate Change 2: 619–622.

Ellison AE, Degrassi AL. 2017. All species are important, but some species are more important than others. Journal of Vegetation Science 28: 669–671.

Foden WB, Butchart SHM, Stuart SN, Vie’ J-C, Akcakaya HR, et al. 2013. Identifying the world’s most climate change vulnerable species: a systematic trait-based assessment of all birds, amphibians and corals. PLoS ONE 8: e65427.

Foden WB, Young BE Akcakaya HR, Garcia RA, Hoffmann AA, et al. 2018. Climate change vulnerability assessment of species. Wiley Interdisciplinary Reviews Climate Change 10: e551

Fordham DA, Resit Akçakaya H, Araújo MB, Elith J, Keith DA, Pearson R, Auld TD, Mellin C, Morgan JW, Regan TJ, Tozer M, Watts MJ, White M, Wintle BA, Yates C, Brook BW. 2012. Plant extinction risk under climate change: are forecast range shifts alone a good indicator of species vulnerability to global warming? Global Change Biology 18: 1357–1371.

Freeman BG, Lee-Yaw JA, Sunday JM, Hargreaves AL (2018) Expanding, shifting and shrinking: the impact of global warming on species’ elevational distributions. Global Ecology and Biogeography 27: 1268–1276.

Gallagher RV, Allen S, Wright IJ. 2019. Safety margins and adaptive capacity of vegetation to climate change. Scientific Reports 9: 8241.

Gaston KJ. 2011. Common ecology. BioScience 61: 354–362.

Gaston KJ, Fuller RA. 2007. Commonness, population depletion and conservation biology. Trends in Ecology and Evolution 23: 14–19.

Geiser F, Broome LS. 1991. Hibernation in the mountain pygmy possum *Burramys parvus* (Marsupialia). Journal of Zoology 223: 593–602.

Geyer J, Kiefer I, Kreft S, Chavez V, Salafsky N, et al. 2011. Classification of climate-change-induced stresses on biological diversity. Conservation Biology 25: 708–715.

Gibson-Reinemer DK, Rahel FJ. 2015. Inconsistent range shifts within species highlight idiosyncratic responses to climate warming. PLoS ONE 10: e0132103.

Gigerenzer G, Edwards A. 2003. Simple tools for understanding risks: from innumeracy to insight. BMJ: British Medical Journal 327: 741–744.

Graae BJ, Ejrnæs R, Lang SI, Meineri E, Ibarra PT, Bruun HH. 2011. Strong microsite control of seedling recruitment in tundra. Oecologia 166: 565–576.

Grabherr G, Gottfried M, Pauli H. 1994. Climate effects on mountain plants. Nature 369: 448.

Granger Morgan M, Pitelka LF, Shevliakova E. 2001. Elicitation of expert judgments of climate change impacts on forest ecosystems. Climatic Change 49: 279–307.

Green K, Osborne W. 1994. Wildlife of the Australian Snow-Country. Reed Press, Sydney.

Green K, Slatyer R. 2020. Arthropod community composition along snowmelt gradients in snowbeds in the Snowy Mountains of south-eastern Australia. Austral Ecology 45: 144–157.

Green K, Stein JA. 2015. Modeling the thermal zones and biodiversity on the high mountains of Meganesia: the importance of local differences. Arctic, Antarctic, and Alpine Research 47: 671–680.

Griffin PC, Hoffmann AA. 2012. Mortality of Australian alpine grasses *(Poa* spp.) after drought: species differences and ecological patterns. Journal of Plant Ecology 5: 121–133.

Guisan A, Thuiller W. 2005. Predicting species distribution: offering more than simple habitat models. Ecology Letters 8: 993–1009.

Halloy SRP, Mark AF. 2003. Climate-change effects on alpine plant biodiversity: a New Zealand perspective on quantifying the threat. Arctic, Antarctic, and Alpine Research 35: 248–254.

Hanea A, McBride M, Burgman M, Wintle B, Fidler F, Flander L Twardy, CR, Manning B, Mascaro S. 2016. Investigate Discuss Estimate Aggregate for structured expert judgement. International Journal of Forecasting 33: 267–269.

Hanea AM, McBride MF, Burgman MA, Wintle BC. 2018. The value of performance weights and discussion in aggregated expert judgments. Risk Analysis 38: 1781–1794

Hargreaves AL, Samis KE, Eckert CG. 2014. Are species’ range limits simply niche limits writ large? A review of transplant experiments beyond the range. The American Naturalist 183: 157–173.

Hemming V, Burgman MA, Hanea AM, McBride MF, Wintle BC. 2018. A practical guide to structured expert elicitation using the IDEA protocol. Methods in Ecology and Evolution 9: 169–181.

HilleRisLambers J, Harsch MA, Ettinger AK, Ford KR, Theobald EJ. 2013. How will biotic interactions influence climate change–induced range shifts? Annals of the New York Academy of Sciences 1297: 112–125.

Hoffmann AA, Chown SL, Clusella-Trullas S. 2013. Upper thermal limits in terrestrial ectotherms: how constrained are they? Functional Ecology 27: 934–949.

Hoffmann AA, Rymer PD, Byrne M, Ruthrof KX, Whinam J, McGeoc M, Bergstrom DM, Guerin GR, Sparrow B, Joseph L, Hill SJ, Andrew NR, Camac J, Bell N, Riegler M, Gardner JL, Williams SE. 2019. Impacts of recent climate change on terrestrial flora and fauna: Some emerging Australian examples. Austral Ecology 44: 3–27

Holling C. 1973. Resilience and stability of ecological systems. Annual Review of Ecology and Systematics 4: 1–23.

IPCC 2014. Glossary. Intergovernmental Panel on Climate Change.

Kirkpatrick JB, Bridle KL. 1999. Environment and floristics of ten Australian alpine vegetation formations. Australian Journal of Botany 47: 1–21.

Kobiv Y. 2017. Response of rare alpine plant species to climate change in the Ukrainian Carpathians. Folia Geobotania 52: 217–226.

Krueger T, Page T, Hubacek K. Smith L, Hiscock K. 2012. The role of expert opinion in environmental modelling. Environmental Modelling & Software 36: 4–18.

La Sorte FA, Jetz W. 2010. Projected range contractions of montane biodiversity under global warming. Proceedings of the Royal Society B: Biological Sciences 277: 3401–3410.

Lawler JJ, Shafer S. L, White D, Kareiva P, Maurer EP, Blaustein AR, Bartlein PJ. 2009. Projected climate-induced faunal change in the Western Hemisphere. Ecology 90: 588–597.

Lenoir J, Gegout JC, Marquet PA, De Ruffray P, Brisse H. 2008. A significant upward shift in plant species optimum elevation during the 20th century. Science 320: 1768–1771.

Lenoir J, Gégout J.-C, Guisan A, Vittoz P, Wohlgemuth T, Zimmermann NE, et al. 2010. Going against the flow: potential mechanisms for unexpected downslope range shifts in a warming climate. Ecography 33: 295–303.

Louthan AM, Doak DF, Angert AL. 2015. Where and when do species interactions set range limits? Trends in Ecology and Evolution 30: 780–792.

Low Choy S, O’Leary R, Mengersen K. 2009. Elicitation by design in ecology: using expert opinion to inform priors for Bayesian statistical models. Ecology 90: 265–277.

Martin TG, Burgman MA, Fidler F, Kuhnert PM, Low-Choy S, McBride M, Mengersen K. 2012. Eliciting expert knowledge in conservation science. Conservation Biology 26: 29–38.

McGowan H, Callow JN, Soderholm J, McGrath G, Campbell M, Zhao J-X. 2018. Global warming in the context of 2000 years of Australian alpine temperature and snow cover. Scientific Reports 8: 4394.

Michalet R, Schöb C, Lortie CJ, Brooker RW, Callaway RM. 2014. Partitioning net interactions among plants along altitudinal gradients to study community responses to climate change. Functional Ecology 28: 75–86.

Millenium Ecosystem Assessment 2005. Ecosystems and Human Well-Being: Synthesis. Island Press, Washington.

Morgan JW, Dwyer JM, Price JN, Prober SM, Power SA, Firn J, Moore JL, Wardle GM, Seabloom EW, Borer ET, Camac JS. 2016. Species origin affects the rate of response to inter-annual growing season precipitation and nutrient addition in four Australian native grasslands. Journal of Vegetation Science 27: 1164–1176.

Morgan JW, Venn SE. 2017. Alpine plant species have limited capacity for long-distance seed dispersal, Plant Ecology 218: 813–819.

Nolan RH, Boer MM, Collins L, Resco de Dios V, Clarke H, Jenkins M, Kenny B, Bradstock RA. 2020. Causes and consequences of eastern Australia’s 2019–20 season of mega-fires. Global Change Biology 26: 1039–1041.

Normand S, Zimmermann NE, Schurr FM, Lischke H. 2014. Demography as the basis for understanding and predicting range dynamics. Ecography 37: 1149–1154.

Ofori BY, Stow AJ, Baumgartner JB, Beaumont LJ. 2017. Influence of adaptive capacity on the outcome of climate change vulnerability assessment. Scientific Reports 7: 12979.

Oppenheimer M, Oreskes N, Jamieson D, Brysse K, O’Reilly J, Shindell M, Wazek M. 2019. Discerning Experts: The Practices of Scientific Assessment for Environmental Policy. University of Chicago Press, Chicago.

Pauli H, Gottfried M, Reiter K, Klettner C, Grabherr G. 2007. Signals of range expansions and contractions of vascular plants in the high Alps: observations (1994–2004) at the GLORIA master site Schrankogel, Tyrol, Austria. Global Change Biology 13: 147–156.

Rumpf SB, Hülber K, Zimmermann NE, Dullinger S. 2019. Elevational rear edges shifted at least as much as leading edges over the last century. Global Ecology and Biogeography 28: 533–543.

Sanchez-Bayo F, Green K. 2013. Australian snowpack disappearing under the influence of global warming and solar activity. Arctic, Antarctic, and Alpine Research 45: 107–118.

Silverstein RN, Correa AM, Baker AC. 2012. Specificity is rarely absolute in coral-algal symbiosis: implications for coral response to climate change. Proceedings of the Royal Society B. 279: 2609–2618.

Smith MD, Knapp AK. 2003. Dominant species maintain ecosystem function with non-random species loss. Ecology Letters 6: 509–517.

Smith MD, Koerner SE, Knapp AK, Avolio ML, Chaves FA, Denton EM, Dietrich J, Gibson DJ, Gray J, Hoffman AM, Hoover DL, Komatsu KJ, Silletti A, Wilcox KR, Yu Q, Blair JM (2020) Mass ratio effects underlie ecosystem responses to environmental change. Journal of Ecology 108: 855–864.

Speirs-Bridge A, Fidler F, McBride M, Flander L, Cumming G, Burgman M. 2010. Reducing overconfidence in the interval judgments of experts. Risk Analysis 30: 512–523.

Steinbauer MJ, et al. 2018. Accelerated increase in plant species richness on mountain summits is linked to warming. Nature 556: 231.

Steinbauer K, Lamprecht A, Semenchuk P, Winkler M, Pauli H. 2020. Dieback and expansions: species-specific responses during 20 years of amplified warming in the high Alps. Alpine Botany 130: 1–11.

Svenning J-C, Sandel B. 2013. Disequilibrium vegetation dynamics under future climate change. American Journal of Botany 100: 1266–1286.

Tingley MW, Koo MS, Moritz C, Rush AC, Beissinger SR. 2012. The push and pull of climate change causes heterogeneous shifts in avian elevational ranges. Global Change Biology 18: 3279–3290.

Umbers KDL, Mappes J. 2015. Postattack deimatic display in the mountain katydid, Acripeza reticulata, Animal Behaviour 100: 68–73.

Venn S, Kirkpatrick JB, McDougall K, Walsh N, Whinam J, Williams RJ. 2017. Alpine, sub-alpine and sub-Antarctic vegetation of Australia. In: D.A. Keith (ed.), Australian Vegetation. pp. 461–490. Cambridge University Press, Cambridge.

Warrant E, Frost B, Green K, Mouritsen H, Dreyer D, Adden A, Brauburger K, Heinze S. 2016. The Australian Bogong Moth *Agrotis infusa*: a long-distance nocturnal navigator. Frontiers in Behavioral Neuroscience 10: 77. doi: 10.3389/fnbeh.2016.00077

Wahren C-H, Camac JS, Jarrad FC, Williams RJ, Papst WA, Hoffmann AA. 2013. Experimental warming and long-term vegetation dynamics in an alpine heathland. Australian Journal of Botany 61: 36–51.

Williams RJ, McDougall KL, Wahren C.-H, Rosengren NJ, Papst WA. 2006. Alpine landscapes. In: Ecology: an Australian Perspective (eds. P. M. Attiwill & B. Wilson), pp. 557–72. Oxford University Press, Oxford.

Williams RJ, Papst WA, McDougall KL, et al. 2014. Alpine ecosystems. In: Biodiversity and Environmental Change: Monitoring, Challenges and Directions (eds. D. Lindenmayer, E. Burns, N. Thurgate & A. Lowe), pp. 167–212. CSIRO Publishing, Melbourne.

Williams RJ, Wahren C.-H, Stott KAJ, Camac JS, White M, Burns E, Harris S, Nash M, Morgan JW, Venn S, Papst WA, Hoffmann AA. 2015. An International Union for the Conservation of Nature Red List ecosystems risk assessment for alpine snow patch herbfields, south-eastern Australia. Austral Ecology 40: 433–443.

Wilson RJ, Gutiérrez D, Gutiérrez J, Martínez D, Agudo R, Monserrat VJ. 2005. Changes to the elevational limits and extent of species ranges associated with climate change. Ecology Letters 8: 1138–1146.

Wintle BC, Fidler F, Vesk PA, Moore JL. 2012. Improving visual estimation through active feedback. Methods in Ecology and Evolution 4: 53–62.

Wipf S, Stoeckli V, Bebi P. 2009. Winter climate change in alpine tundra: plant responses to changes in snow depth and snowmelt timing. Climatic Change 94: 105–121.

Zylstra PJ. 2018. Flammability dynamics in the Australian Alps. Austral Ecology 43: 578–591.

