## Supplementary Material for "Predicting species and community responses to global change in Australian mountain ecosystems using structured expert judgement"

### Supplemental Material


Supplemental methods 2
  Pre elicitation 2
    Expert selection 2
      Table S1 - Experts who contributed to the expert elicitation workshops 4
    Survey questions 6
      Table S2 - Survey provided to participants before the workshops 7
    Calibration dataset 8
    Climate scenario 9
      Table S3 - climate scenario used in the animal elicitation workshop. 9
    Target species 10
  Elicitation - data collection 15
    Initial estimates 15
    Discussion of initial estimates and submission of revised estimates 15
  Post-elicitation - data analysis 16
    Aggregation of expert judgements: weighted vs equal weighting 16
  Compilation of plant trait and environment data 17
Supplemental Figures 19
      Figure S1 – Schematic of IDEA protocol 20
      Figure S2 – Adaptive capacity & plant environmental attributes 21
      Figure S3 – Adaptive capacity & continuous plant species traits 22
      Figure S4 – Adaptive capacity & plant species categorical traits 23
Data link 24
References 24

### Supplemental methods

The plant and animal expert elicitation projects were undertaken in July 2017 and November 2018, respectively. The plant workshop was conducted at La Trobe University, Melbourne and focused on estimating current and future cover of Australian alpine plants and plant communities. The animal workshop was hosted by Western Sydney University and conducted at Katoomba, NSW and focused on estimating current and future abundance and distribution patterns. In both cases, participants provided written consent to take part in the study.

### Pre elicitation

#### Expert selection

Experts ($n$ = 22 for plants, $n$ = 17 for animals, $n$ = 2 shared between workshops; Table S1) were selected to represent the breadth of expertise in alpine botany, zoology and ecology in Australia. Authors compiled an initial list and, to avoid bias, asked the initially identified experts to nominate other experts who they thought should be included. The groups were, we believe, a representative sample of the breadth of experience from what is a relatively small total number of relevant subject experts. Qualitative data were first collected from all experts to gain insight into their *a priori* judgement on the definitions and drivers of adaptive capacity prior to the elicitation workshop (Table S2).

The list of experts included a) academic researchers and post-graduate students actively involved in botanical, zoological and ecological research in the alps, b) management agency staff involved in field ecology, surveys and management of the alps, c) staff from botanical gardens, zoos and museums in representative states with extensive experience in the alps. Ultimately the participants in the workshop included 9 women and 13 men in the plant group and 8 women and 9 men in the animal group.  In total, our experts had ~500 hundred years' experience with Australian mountain plants and ~200 years with mountain animals. Individual experience ranged from 5 to 50 years for plant experts and 1 to 40 years for animal experts. All experts had experience working in one or more of the major mountain regions within the alps, which can be loosely delineated by state (Tasmania, Victoria, New South Wales, Australian Capital Territory).

Training was provided to all experts prior to the workshops. The information provided included an explanation of the IDEA protocol and the question format used for the exercise. For the plant workshop, the training was provided via an excel sheet with supporting documents. For the animal workshop, we gave a two-hour webinar training session in which experts were guided through an example that demonstrated how to answer the questions. For both groups, we emphasised to the experts the important principles necessary to maximise the data quality before they filled out the first survey. This included following the same order when estimating the lower and upper bounds before the best estimate, providing their confidence in their answer, and ensuring all estimates were completed before subsequent group discussion so that they 'anchor' to their own answers (Hanea et al. 2016). Experts were also given a brief refresher training session on how to answer the questions in person at the workshops and prior to each discussion of the calibration or survey answers respectively.

**Table S1 - Experts who contributed to the expert elicitation workshops**

The names and affiliations of all the participants who both attended and provided estimates at one or both workshops. Note that two participants opted to not to be named.

| Name | Affiliation | Workshop Participation |
|---|---|---|
| Emma Burns | Australian National University | Plant |
| James Camac | The University of Melbourne | Plant |
| Michael Driessen | University of Tasmania | Animal |
| Michael Doherty | Australian National University | Plant |
| Francisco Encinas-Viso | CSIRO | Animal |
| Sonya Geange | Australian National University | Plant & Animal |
| Louise Gilfedder | University of Tasmania | Plant |
| Lydia Guja | Australian National Botanic Gardens | Plant |
| Margaret Haines | Museums Victoria | Animal |
| Ary Hoffmann | The University of Melbourne | Plant & Animal |
| Geoff Hope | Australian National University | Plant |
| David Keith | University of New South Wales | Plant |
| Casey Kirchhoff | University of New South Wales | Animal |
| Bryan Lessard | CSIRO | Animal |
| Joe McAuliffe | Desert Botanical Garden | Plant |
| Keith McDougall | Department of Planning, Industry and Environment (NSW) | Plant |
| Scott Mooney | University of New South Wales | Plant |
| John Morgan | La Trobe University | Plant |
| Giselle Muschett | Pontificia Universidad Catolica de | Animal |

| | | |
|---|---|---|
| Julia Mynott | La Trobe University | Animal |
| Michael Nash | La Trobe University | Animal |
| Adrienne Nicotra | Australian National University | Plant |
| Juanita Rodriguez | CSIRO | Animal |
| Ben Scheele | Australian National University | Animal |
| James Shannon | La Trobe University | Plant |
| Rachel Slatyer | Australian National University | Animal |
| Phil Suter | La Trobe University | Animal |
| Kate Umbers | Western Sydney University | Plant |
| Susanna Venn | Deakin University | Plant |
| Peter Vesk | The University of Melbourne | Plant |
| Neville Walsh | Royal Botanic Gardens Melbourne | Plant |
| Erik Wapstra | University of Tasmania | Animal |
| Geoff While | University of Tasmania | Animal |
| Richard Williams | Charles Darwin University | Plant |
| Jennie Winham | University of Tasmania | Plant |
| Genevieve Wright | University of Sydney | Plant |

#### Survey questions

The survey questions were developed in advance with consultation among the workshop organisers, discussed at length with participants at the start of each workshop, and then refined to avoid any biological and linguistic ambiguities. The final versions of the questions are presented below (Table S2). Because there is no accepted method by which to quantify or compare adaptive capacity across plants and animals, we developed questions based on estimates of percent cover for plants and for abundance and elevation range for animals for the present day (2017 and 2018, respectively) and in 2050. Adaptive capacity reflects the potential of a species to maintain or increase its abundance in the face of climate change (Dawson et al. 2011; Ofori et al. 2017). Therefore, for plants, we first asked our experts to visualise the percent cover for a given species in an average sampling plot (10 x 10 m) for a given plant community type. For the animals, the size of the sample area was adjusted as appropriate to each organism and ranged from 1 $m^2$ for small, high-density animals like the bogong moth (*Agrotis infusa*) to 5000 $m^2$ for large, mobile animals like the broad-toothed rat (*Mastacomys fuscus*) and alpine copperhead snake (*Austrelaps ramsayi*). We examined these estimated values relative to the estimates of cover in 2050 as an indication of future performance, and thus adaptive capacity.

For the plants, answers were obtained using a 4-point elicitation approach where experts were asked to estimate their (1) lowest plausible value, (2) highest plausible value, (3) best estimate, and (4) their confidence that the truth falls within their lower and upper limit. We subsequently concluded that the confidence estimates were misinterpreted and thus, unreliable. As a consequence, the animal elicitation workshop utilised a 3-point elicitation approach such that experts were only asked to provide their lowest plausible estimate as a $5^{th}$ percentile, their highest plausible estimate as the $95^{th}$ percentile and their best estimate as the $50^{th}$ percentile of values (i.e. their best estimate).

**Table S2 - Survey provided to participants before the workshops**

Survey tools (questions) used in the plant and animal elicitation process.

**Background information and qualitative judgements asked prior to both the plant and animal elicitations**

1. Please describe your experience working in the Australian mountains, including number of years.
2. Please describe your experience working with alpine plants/animals, including number of years, locations where you've worked, focal taxa (animal only).
3. How would you describe yourself professionally (this might include your field and your most relevant (current or previous) employment)?
4. What is your interpretation of the following terms? (at most 100 words each)
    a. adaptive capacity of a species
    b. functional importance of a species
5. What factors do you think contribute to /influence the adaptive capacity of a species?
6. What characteristics would you need to know about a species to estimate its adaptive capacity?

**Calibration questions (plant elicitation only)**

1. What is the current total elevation range (in metres) for this species, in the indicated vegetation type on the mainland?
2. What would the percent cover of this species, averaged across typical 10m*10m vegetation plots, have been 25 years ago in the vegetation type listed below? The bounds in question 2 refer to lowest and highest plausible values for the percent cover.

**Plant elicitation survey questions**

1. What is the current percent cover of this species, if it were averaged across a series of typical 10m*10m vegetation plots, in which the species occurs, in the vegetation type listed below?
2. What will the future percent cover of this species be, if it were averaged across a series of typical 10m*10m vegetation plot, in the vegetation type listed below, at 2050 (assuming it was present in 2017)?

**Animal elicitation survey questions**

1. Low-elevation Range Limits
    a. What is the low-elevation range limit (in metres above sea level) of this species?
    b. Climate conditions are projected to change as outlined in the table below. Given this scenario, what will the low-elevation range limit (in meters above sea level) of this species be in 2050?
2. High-elevation Range Limits
    a. What is the high-elevation range limit (in metres above sea level) of this species?

b. Climate conditions are projected to change as outlined in the table below. Given this scenario, what will the high-elevation range limit (in metres above sea level) of this species be in 2050?

3. Abundance
   a. When this species is present, how many individuals are there, on average, at their estimated peak abundance in the habitat type and area specified?
   b. Climate conditions are projected to change as outlined in the table below. Given this scenario, by 2050, how many individuals will there be in the same location as you envisaged for Q3A, on average, at their peak abundance?

#### Calibration dataset

A calibration dataset was compiled for the plant workshop. Calibration questions are designed to measure the calibration and informativeness of the expert's responses to questions for which data are available and the answer is known by the analyst, but not directly available to the expert at the time of the elicitation. Each expert was asked to estimate the elevation ranges of selected plant species in the Australian Alps in 2017. In addition to this, we also asked experts to estimate what the average percent cover of each species would have been 25 years ago in a typical 10 x 10 m vegetation plot in which the species occurs in the plant community type most associated with the species. A 10 x 10 m plot was chosen as a typical plot size for vegetation sampling in plant communities with small stature. We used percent cover in a typical vegetation plot as an indicator of the dominance of the species in the given vegetation community. The species included in the plant calibration exercise were a subset of the total species pool used in the main surveys. The elevation ranges were determined using floristic plot data (1682 plots) derived from McDougall and Walsh (2007). Mean cover for species in particular communities was calculated as follows: plots were divided into communities according to the classification of McDougall and Walsh (2007); random numbers were generated within the cover range of the values used in the surveys (on the Braun-Blanquet cover scale); mean cover values were then calculated for the selected species only for plots in which they occurred.

In the plant exercise, calibration questions were included but subsequent analyses demonstrated that calibration weighted estimates did not differ significantly from the unweighted estimates (see below). As such, calibration weighted estimates were not used in this study, nor applied in the animal elicitation.

#### Climate scenario

In the plant elicitation, several climate change models were discussed and a general consensus was reached that there would be increases in temperature, decreasing precipitation (and less of that falling as snow, and fewer days of snow cover), and increased chance of fire. The plant elicitation explicitly focussed on responses of the species on the mainland (excluding the Tasmanian portion of the distribution of any species that were also found there).

For the animal workshop, we sought to reduce the amount of time spent in discussion of potential climate scenarios and maximise the clarity of questions by providing a specific climate scenario for the year 2050 (Table S3). Trends and numbers were gathered from two technical reports to represent conditions in 2050 relative to 1990 (CSIRO and Bureau of Meteorology 2015). The climate scenarios were provided as a guide to ensure that the experts were thinking about the same climate conditions when making estimates. Experts were asked to include notes on particular aspects of climate that were considered when making their estimates.

**Table S3 - climate scenario used in the animal elicitation workshop.**

Climate scenario for the alpine animal elicitation. Projected increases are coloured orange and projected decreases are coloured purple.

| | Climatic Variable | Mainland Australia | Tasmania |
|---|---|---|---|
| **Temperature** | Mean temperature | ↑ 1.8°C | ↑ 1.1°C |
| **Rain & moisture** | Cool-season rainfall | ↓ 15 % | ↑ 5 % |
| | Hot-season rainfall | ↓ | ↓ |
| | Mean soil moisture | ↓ | ↓ |
| | Frequency of drought | ↑ | -- |
| **Snow** | Area of snow cover for at least 30 days | ↓ 60 % | ↓ 60 % (§ - see footnotes in ref tab) |
| | Area of snow cover for at least 60 days | ↓ 65 % | ↓ 65 % (§ - see footnotes in ref tab) |
| | Days of > 1 cm snow cover | ↓ 50 days | ↓ 50 days (§ - see footnotes in ref tab) |
| | Maximum snow depth | ↓ 50 % | ↓ 50 % (§ - see footnotes in ref tab) |
| | Elevation of snow line | ↑ 270 m (# - see footnotes in ref tab) | ↑ 165 m (# - see footnotes in ref tab) |
| **Fire** | Days with high fire danger | ↑ | ↑ |

[§]No predictions available for Tasmania, but changes are considered to be similar to those on the mainland (Grose et al. 2015).
[#] Based on a predicted increase in the elevation of the snowline of 150m/1°C rise in temperature (Abegg et al. 2007)

#### Target species

In the plant workshop, experts estimated the current (2017) and the 2050 cover of 60 plant species (Table S4A). In addition, experts estimated the 2050 landscape cover of nine alpine plant communities based on the following baseline covers: Feldmark (0.1%), Snowpatch (1%), Grassland/Herbfield (25%), Woodland (24%), Heathland (35%), Bog (5%), Fen (4%), Wet tussock grassland (6%). We selected species to represent a range of abundance and cover in the respective vegetation community.

Twenty-nine animal species were selected for assessment, representing a range of taxonomic and functional groups (Table S4B). The selection included species who were not functionally extinct in the wild and whose distributions extend beyond the alps, but are predominantly found in the alpine or subalpine ecosystems. The species included arthropods (insects and crustaceans) and chordates (fish, amphibians, reptiles and mammals) and one platyhelminth, from both aquatic and terrestrial habitats. The number of experts that provided answers varied for each of the animal species. Three additional species (*Venatrix funesta*, *Notoscolex montiskosciuskoi* & *Percolestus blackburni*) were also originally assessed at the animal workshop, however as fewer than 4 experts provided estimates for these species they excluded from our analyses.

**Table S4.** Species considered in the A) plant and B) animal elicitation workshops, with associated ID, family, community type they typically occur in, and growth form/taxon class.

*A) Plant species*

| *ID* | *Species* | Family | Community | Growth form |
|---|---|---|---|---|
| *1* | *Stylidium montanum* | Stylidiaceae | Woodland | Forb |
| *2* | *Tasmannia xerophila* | Winteraceae | Woodland | Shrub |
| *3* | *Pimelea ligustrina* | Thymeleaceae | Woodland | Shrub |
| *4* | *Eucalyptus pauciflora* | Myrtaceae | Woodland | Tree |
| *5* | *Picris angustifolius* | Asteraceae | Woodland | Forb |
| *6* | *Goodenia hederacea* | Goodeniaceae | Woodland | Forb |
| *7* | *Dianella tasmanica* | Liliaceae | Woodland | Forb |
| *8* | *Oxylobium ellipticum* | Fabaceae | Woodland | Shrub |
| *9* | *Asperula gunnii* | Rubiaceae | Woodland | Forb |
| *10* | *Acaena novae-zelandiae* | Rosaceae | Woodland | Forb |
| *11* | *Podocarpus lawrencei* | Podocarpaceae | Heathland | Shrub |
| *12* | *Pentachondra pumilio* | Epacridaceae | Heathland | Shrub |
| *13* | *Trisetum spicatum* | Poaceae | Heathland | Graminoid |
| *14* | *Pimelea axiflora* | Thymeleaceae | Heathland | Shrub |
| *15* | *Grevillea australis* | Proteaceae | Heathland | Shrub |
| *16* | *Orites lancifolia* | Epacridaceae | Heathland | Shrub |
| *17* | *Acrothamnus montanus* | Epacridaceae | Heathland | Shrub |
| *18* | *Olearia brevipedunculata* | Asteraceae | Heathland | Shrub |
| *19* | *Viola betonicifolia* | Violaceae | Heathland | Forb |

| | | | | |
|---|---|---|---|---|
| *20* | *Kunzea muelleri* | Myrtaceae | Heathland | Shrub |
| *21* | *Prostanthera cuneata* | Lamiaceae | Heathland | Shrub |
| *22* | *Hovea montana* | Fabaceae | Heathland | Shrub |
| *23* | *Celmisia pugioniformis* | Asteraceae | Grassland | Forb |
| *24* | *Podolepis robusta* | Asteraceae | Grassland | Forb |
| *25* | *Gentianella muelleriana* | Gentianaceae | Grassland | Forb |
| *26* | *Aciphylla glacialis* | Apiaceae | Grassland | Forb |
| *27* | *Poa hiemata* | Poaceae | Grassland | Graminoid |
| *28* | *Craspedia aurantia* | Asteraceae | Grassland | Forb |
| *29* | *Scleranthus biflorus* | Caryophyllaceae | Grassland | Forb |
| *30* | *Prasophyllum alpestre* | Orchidaceae | Grassland | Forb |
| *31* | *Poa costiniana* | Poaceae | Grassland | Graminoid |
| *31* | *Poa costiniana* | Poaceae | Wetland | Graminoid |
| *32* | *Oreomyrrhis eriopoda* | Apiaceae | Grassland | Forb |
| *33* | *Wahlenbergia ceracea* | Campanulaceae | Grassland | Forb |
| *34* | *Leptorhynchos squamatus* | Asteraceae | Grassland | Forb |
| *35* | *Carex breviculmis* | Cyperaceae | Grassland | Graminoid |
| *36* | *Senecio gunnii* | Asteraceae | Grassland | Forb |
| *37* | *Agrostis venusta* | Poaceae | Grassland | Graminoid |
| *38* | *Sphagnum cristatum* | Sphagnaceae | Wetland | Moss |
| *39* | *Carpha alpina* | Cyperaceae | Wetland | Graminoid |
| *40* | *Richea continentis* | Epacridaceae | Wetland | Shrub |
| *41* | *Oreomyrrhis ciliata* | Apiaceae | Wetland | Forb |
| *42* | *Drosera arcturi* | Droseraceae | Wetland | Forb |
| *43* | *Carex gaudichaudina* | Cyperaceae | Wetland | Graminoid |
| *44* | *Oreobolus distichus* | Cyperaceae | Wetland | Graminoid |
| *45* | *Astelia alpina* | Liliaceae | Wetland | Forb |
| *46* | *Epacris petrophila* | Epacridaceae | Wetland | Shrub |
| *47* | *Epacris paludosa* | Epacridaceae | Wetland | Shrub |
| *48* | *Empodisma minus* | Restionaceae | Wetland | Graminoid |
| *49* | *Baeckea gunniana* | Myrtaceae | Wetland | Shrub |
| *50* | *Montia australasica* | Portulacaceae | Snowpatch | Forb |
| *51* | *Psychrophila introloba* | Ranunculaceae | Snowpatch | Forb |
| *52* | *Oreomyrrhis pulvinifica* | Apiaceae | Snowpatch | Forb |
| *53* | *Luzula acutifolia* | Cyperaceae | Snowpatch | Graminoid |

| | | | | |
|---|---|---|---|---|
| *54* | *Plantago muelleri* | Plantaginaceae | Snowpatch | Forb |
| *55* | *Ewartia nubigena* | Asteraceae | Snowpatch | Forb |
| *56* | *Celmisia costiniana* | Asteraceae | Snowpatch | Forb |
| *57* | *Poa fawcettiae* | Poaceae | Snowpatch | Graminoid |
| *58* | *Carex hebes* | Cyperaceae | Snowpatch | Graminoid |
| *59* | *Agrostis muelleriana* | Poaceae | Snowpatch | Graminoid |
| *60* | *Rytidosperma nudiflorum* | Poaceae | Snowpatch | Graminoid |

*B) Animal species*

| *ID* | *Species* | Family | Water-centric | Taxon |
|---|---|---|---|---|
| *1* | *Agrotis infusa* | Noctuidae | Not-water-centric | Insect |
| *2* | *Pseudemoia cryodroma* | Scincidae | Not-water-centric | Lizard |
| *3* | *Carinascincus microlepidotus* | Scincidae | Not-water-centric | Lizard |
| *4* | *Burramys parvus* | Burramyidae | Not-water-centric | Mammal |
| *5* | *Oncopera alpina* | Hepialidae | Not-water-centric | Insect |
| *6* | *Leioproctus obscurus* | Colletidae | Not-water-centric | Insect |
| *7* | *Mastacomys fuscus* | Muridae | Not-water-centric | Mammal |
| *8* | *Kosciuscola tristis* | Acrididae | Not-water-centric | Insect |
| *9* | *Dirce aesiodora* | Geometridae | Not-water-centric | Insect |
| *10* | *Tasmanalpina clavata* | Acrididae | Not-water-centric | Insect |
| *11* | *Carinascincus greeni* | Scincidae | Not-water-centric | Lizard |
| *12* | *Liopholis guthega* | Scincidae | Not-water-centric | Lizard |
| *13* | *Eulamprus kosciuskoi* | Scincidae | Not-water-centric | Lizard |
| *14* | *Monistria concinna* | Pyrgomorphidae | Not-water-centric | Insect |
| *15* | *Polyzosteria viridissima* | Blattidae | Not-water-centric | Insect |
| *16* | *Acripeza reticulata* | Tettigoniidae | Not-water-centric | Insect |
| *17* | *Rankinia diemensis* | Agamidae | Not-water-centric | Lizard |
| *18* | *Pseudophryne pengilleyi* | Myobatrachidae | Water-centric | Frog |
| *19* | *Galaxias supremus* | Galaxidae | Water-centric | Fish |
| *20* | *Philoria frosti* | Limnodynastidae | Water-centric | Frog |
| *21* | *Archipetalia auriculata* | Austropetaliidae | Water-centric | Insect |
| *22* | *Thaumatoperla alpina* | Eustheniidae | Water-centric | Insect |
| *23* | *Paragalaxius julianus* | Galaxidae | Water-centric | Fish |
| *24* | *Austroaeschna flavomaculata* | Telephlebiidae | Water-centric | Insect |
| *25* | *Euastacus rieki* | Parastacidae | Water-centric | Crustacean |

| | | | | |
|---|---|---|---|---|
| *26* | *Caenoplana coerulea* | Geoplanidae | Water-centric | Worm |
| *27* | *Coloburiscoides giganteus* | Coloburiscidae | Water-centric | Insect |
| *28* | *Crinia nimbus* | Myobatrachidae | Water-centric | Frog |
| *29* | *Anaspides tasmaniae* | Anaspidesidae | Water-centric | Crustacean |

### Elicitation - data collection

#### Initial estimates

The plant expert elicitation process took place in three rounds of 16-25 species each. Initial calibration and a first survey were emailed to participants and conducted by experts individually, prior to the meeting of all experts in the discussion workshop in June 2017. A second survey with additional taxa, as well as an additional survey in which the same questions were applied to the five vegetation communities as a whole, was conducted during the meeting; a final survey was completed remotely three weeks following. For the community survey, experts were supplied with an estimate of the current percentage of alpine/subalpine areas covered by that community type.

Animal experts answered the questions in two main rounds each containing 14 or 15 species and a third and final round that included species added at the workshop. Experts were given two weeks to complete and return the target surveys with their initial estimates, ensuring that all questions were completed by experts independently prior to the discussion workshop. For species added at the workshop, animal experts made their initial estimates at or after the workshop and then participated in discussions solely via video link, or were provided detailed notes on the discussion to revise their initial estimates.

#### Discussion of initial estimates and submission of revised estimates

To facilitate discussion, participants were provided with a figure showing anonymized initial estimates (i.e. best estimate and upper and lower bounds converted to 90% confidence limits) made by members of their group for each species (Fig S1). Experts discussed, in turn, the reasons and logic behind their responses, with particular focus on those questions for which experts' estimates were most divergent, to explore whether different experts had different knowledge or deeper insights. Following these discussions, the experts were asked to return to and adjust their initial judgements if the discussion had convinced them to change anything. These revised estimates were done anonymously, in private. For discussion, experts were divided into two groups of ~12 people structured to spread participants with regard to expertise and seniority. The majority of species were discussed in person but a few species were discussed over video link. The same format for facilitation was followed for all discussions.

### Post-elicitation - data analysis

#### Aggregation of expert judgements: weighted vs equal weighting

Expert-derived data is often aggregated in one of two ways. The first, and simplest approach, is to assume all experts are equally weighted. The alternative is to weight each expert relative to their performance on answering calibration questions (i.e. the questions with known answers). The plant expert elicitation used a calibration question but performance measures (i.e., informativeness and calibration; Cooke 1991) revealed that the two aggregation methods performed similarly. Furthermore, post-elicitation discussions with experts revealed that while best estimates were consistently estimated in a similar fashion across experts, considerable variability existed in how experts interpreted, and thus, estimated their bounds and confidence. For these reasons, our analysis of both the plant and animal data focused on using the individual expert equally weighted best estimates and not their estimated uncertainty defined by their bounds and estimated confidence.

### Compilation of plant trait and environment data

Based on the pre-elictiation survey, we compiled a list of environmental and functional traits that experts nominated as drivers of adaptive capacity for the plant species. We then compiled a database based on these traits. Functional trait data are scarce for Australian alpine animals, and were not included in the analysis.

For each plant species, elevation range, maximum and minimum elevation, area of occupancy and extent of range were determined for each plant species in New South Wales (NSW), Victoria (VIC) and the Australian Capital Territory (ACT). Records were downloaded from Australia's Virtual Herbarium (https://avh.chah.org.au/; accessed: May 2018), cleaned to exclude erroneous data and then elevation (minimum, maximum and range) and Mean Annual Temperature (MAT, minimum, maximum and range) data were extracted for the remaining points. 30 arc second downscaled MAT data were obtained from WorldClim (Hilmans et al 2005). Extent of occurrence (EOO) and area of occupancy (AOO) were calculated using IUCN criteria (Bachman and Moat 2012).

The occurrence data were cleaned in the following way: 1) Distribution records were plotted in ArcMap 10.4 over a topographic map. Then 2) distributions were compared to the expected distribution according to the Flora of NSW (PlantNET: http://plantnet.rbgsyd.nsw.gov.au/) and the Flora of Victoria (vicflora: https://vicflora.rbg.vic.gov.au/). 3) Any points outside the expected distribution were individually examined and deleted if there was no locality information on the herbarium label OR if the locality information on the herbarium layer was inconsistent with the location on the map. 4) Data were inspected for duplicates that have different locations and the one which is not consistent with the label location (or both if necessary) was deleted. 5) Old records (pre-1920) with generic locality info (e.g. Mt Kosciuszko) were deleted as it was not possible to extract accurate location data.

Plant functional trait data were obtained from the experts' published and unpublished data as well as other published and online sources, the flora, and for a few species field specimens were collected to supplement available data.

To determine whether the direction and magnitude of change projected by the experts correlated with the drivers of adaptive capacity that the experts provided in our initial qualitative survey, we calculated a proportional change in cover as a proxy for adaptive capacity. Species for which the experts project a marked decrease in cover over this timeframe have low Adaptive Capacity whereas those that have a large proportional increase in cover have high Adaptive Capacity. We used Equation 1 which yields measures that range from -1 to 1 as our index of Adaptive Capacity (AC):

$$AC = \frac{(Future\ cover - Current\ cover)}{Future\ cover + Current\ cover} \qquad \text{eqn: 1}$$

The values were then plotted against the environmental and trait data based upon the conceptual framework to determine whether the nominated drivers of adaptive capacity were correlated with the average expert's estimates of how taxa would perform over the coming 30+ years.

### Supplemental Figures

**Figure S1 – Schematic of IDEA protocol**

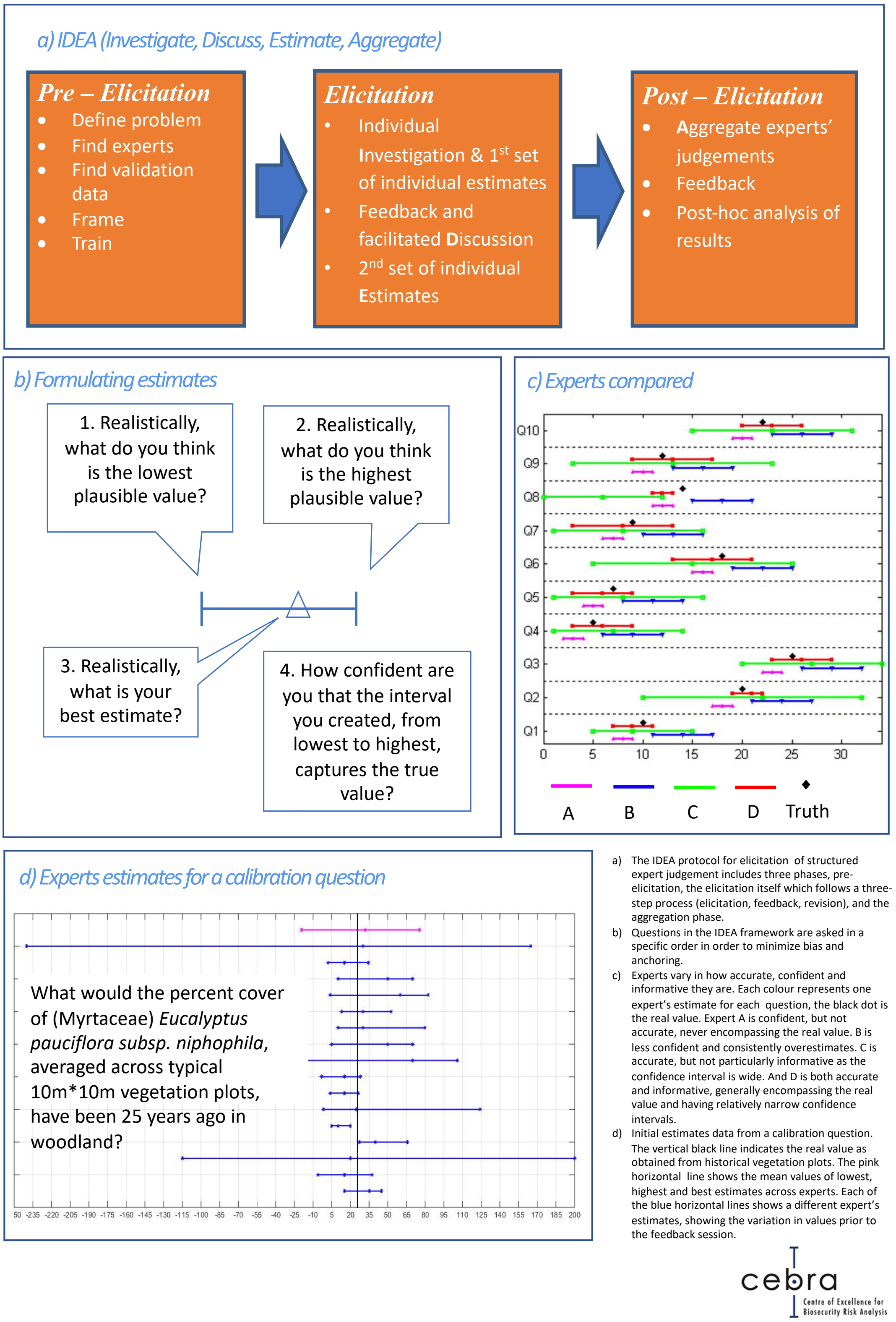


**Fig S1** Method schematic showing the approach used to elicit information from experts and then analyse the resultant data.

**Figure S2 – Adaptive capacity & plant environmental attributes**

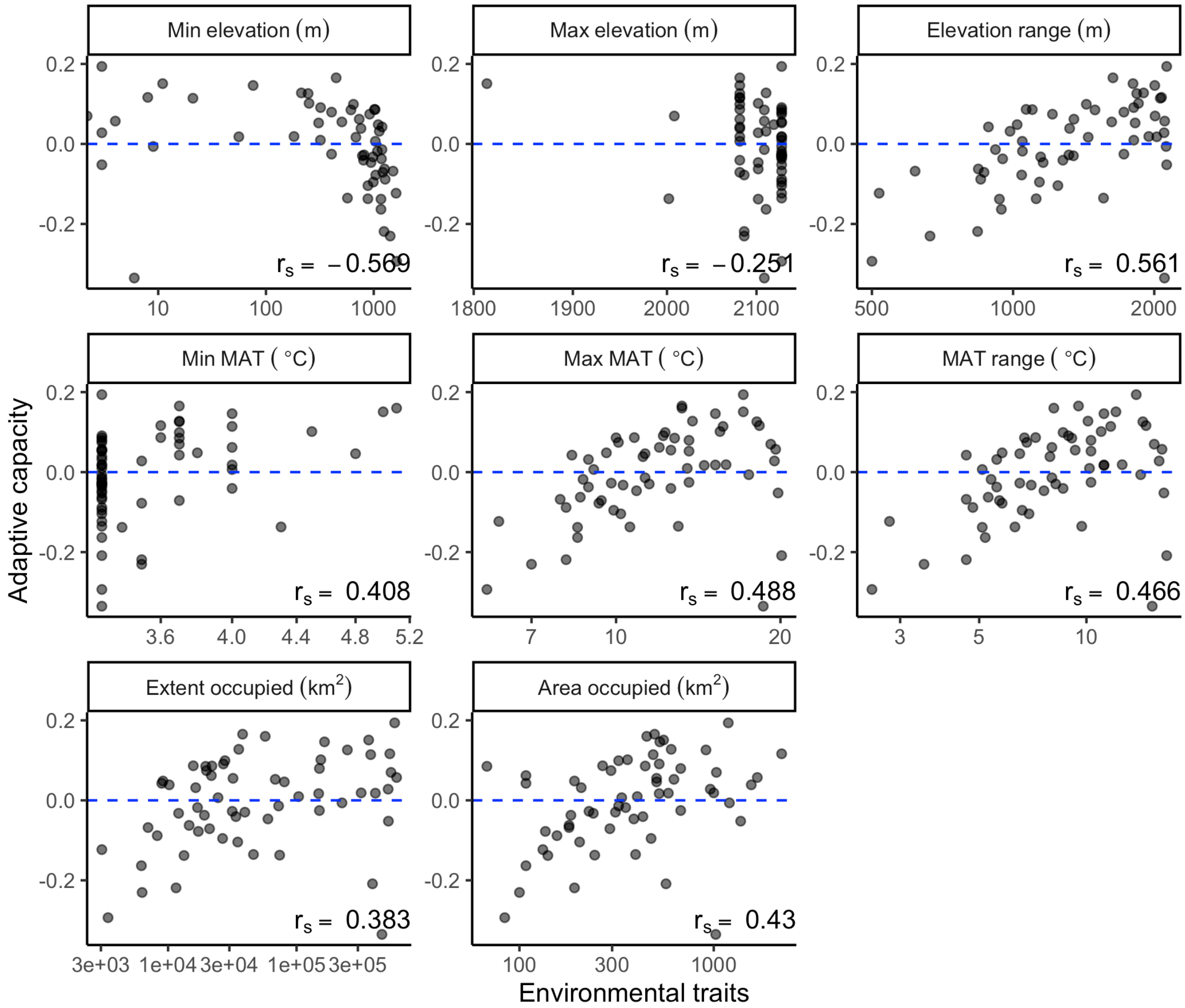


**Fig S2.** Correlations between adaptive capacity and plant species environmental attributes. $r_s$ = Spearman rank correlation. Zero line signifies no expected change in cover between 2017 and 2050.

**Figure S3 – Adaptive capacity & continuous plant species traits**

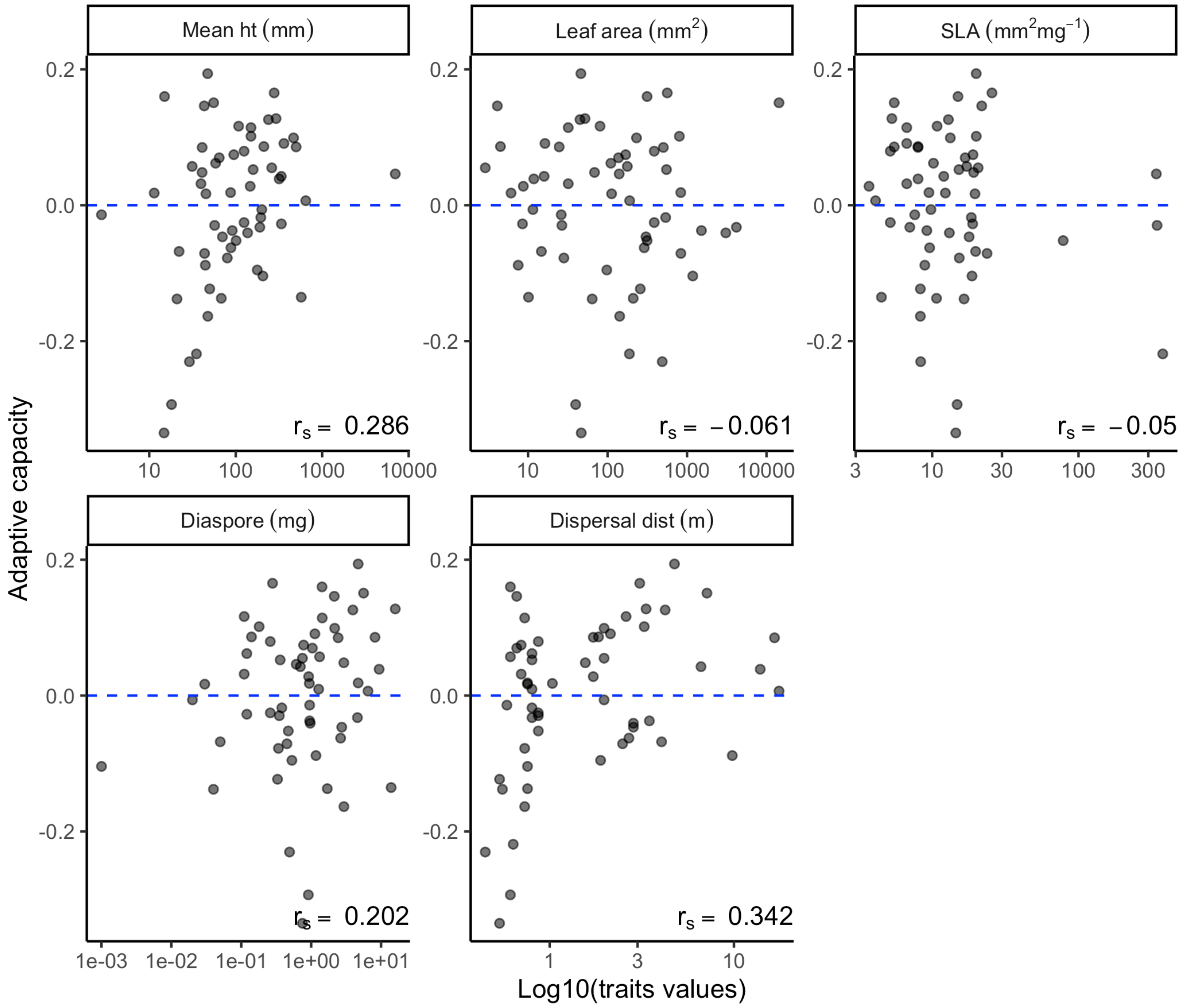


**Fig S3.** Correlations between adaptive capacity and plant species continuous traits. $r_s$ = Spearman rank correlation. Zero line signifies no expected change in cover between 2017 and 2050.

**Figure S4 – Adaptive capacity & plant species categorical traits**

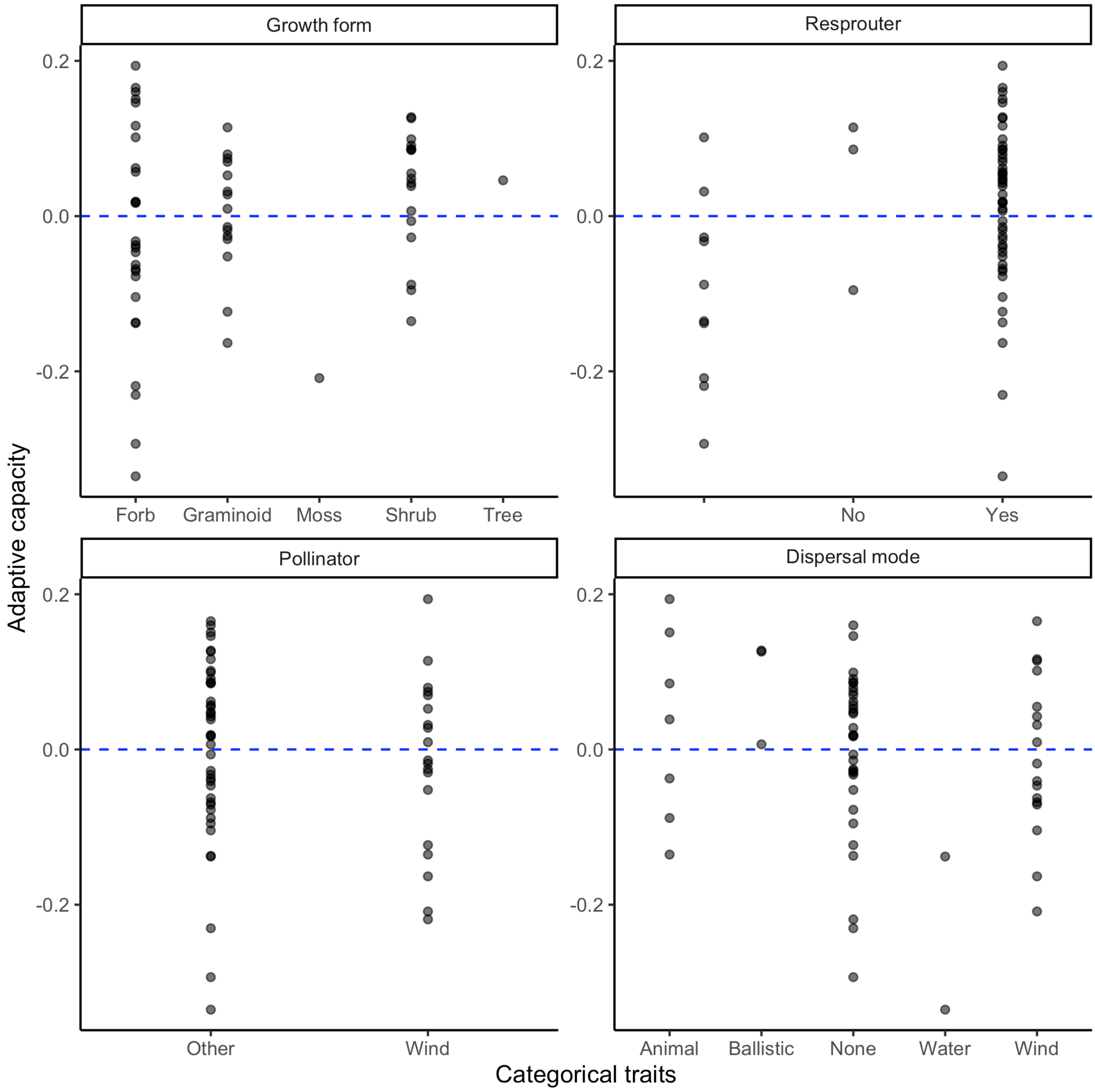


**Fig S4.** Correlations between adaptive capacity and plant species categorical traits. Zero line signifies no expected change in cover between 2017 and 2050.

### Data link

De-identified data and code used to produce figures 1-4 and Supplementary figures S2-S4 can be found at: https://github.com/jscamac/Alpine_Elicitation_Project.

### References


Abegg B, Agrawala S, Crick F, de Montfalcon A. 2007. Climate change impacts and adaptation in winter tourism. In S. Agrawala (ed.) Climate change in the European Alps. Adapting winter tourism and natural hazards management. pp. 25–60. Paris: OECD.

Bachman S, Moat J. 2012. GeoCAT – an open source tool for rapid Red List assessments BGjournal 9: 11-13.

Cooke RM. 1991. Experts in Uncertainty. Oxford, Oxford University Press.

CSIRO and Bureau of Meteorology. 2015. Climate Change in Australia Information for Australia's Natural Resource Management Regions: Technical Report, CSIRO and Bureau of Meteorology, Australia.

Dawson TP, Jackson ST, House JI, Prentice IC, Mace GM. 2011. Beyond predictions: biodiversity conservation in a changing climate. Science **332:** 53–58.

Grose, M. et al. (2015). Southern Slopes Cluster Report, Climate Change in Australia Projections for Australia's Natural Resource Management Regions: Cluster Reports, eds. Ekström, M. et al., CSIRO and Bureau of Meteorology, Australia.

Hanea A, McBride M, Burgman M, Wintle B, Fidler F, Flander L Twardy, CR, Manning B, Mascaro S. 2016. Investigate Discuss Estimate Aggregate for structured expert judgement. International Journal of Forecasting **33:** 267–269.

Hijmans RJ, Cameron SE, Parra JL, Jones PG, Jarvis A. 2005. Very high resolution interpolated climate surfaces for global land areas. International Journal of Climatology **25:** 1965-1978.

Ofori BY, Stow AJ, Baumgartner JB, Beaumont LJ. 2017. Influence of adaptive capacity on the outcome of climate change vulnerability assessment. Scientific Reports **7:** 12979.
